# A contact-mediated transfer mechanism directs *in cis* localization and spread of Xist RNA

**DOI:** 10.64898/2026.10.02.756235

**Authors:** Adam D. Cawte, Guifeng Wei, Flavia Constantinescu, Mafalda Almeida, Heather Coker, Neil Brockdorff

## Abstract

X chromosome inactivation in mammals is driven by the long non-coding RNA Xist, which localizes and spreads across the chromosome from which it is transcribed. Although several factors required for Xist localization have been identified, the underlying mechanism remains unclear. Here, we combine Nuclear-BRITE, a method for visualizing and quantifying the dynamics of individual Xist RNA molecules, with acute depletion of candidate localization factors. We find that Xist molecules remain constrained throughout interphase, consistent with continuous association with chromatin. Building on this observation, we identify a contact-mediated transfer mechanism in which 3D chromosome dynamics promote handover of Xist molecules between chromosomal sites. We further show that the Xist-binding proteins SPEN and HNRNPU play distinct roles in this process by promoting Xist–Xist and Xist–chromatin interactions, respectively. Together, our findings define a mechanism for Xist RNA spreading and provide a framework for understanding its unique chromosome-wide localization.

## Main

Long non-coding RNAs (lncRNAs) play an important role in gene and genome regulation, functioning either *in cis* or in *trans*^1^. A classical model for the former is the *cis*-acting lncRNA Xist, the master regulator of X chromosome inactivation (XCI) in mammals^2,3^. Xist RNA is expressed from the future inactive X chromosome (Xi) in cells of early XX female embryos where it accumulates and spreads *in cis* across the X chromosome. Localized Xist molecules then function by recruiting factors that modify the underlying chromatin and silence X-linked gene expression. A key unresolved question is how Xist RNA retains specificity for the Xi whilst efficiently spreading across the entire X chromosome^4^.

Progress towards understanding localization and spread of Xist RNA has come from several directions. Contemporary studies using genomics approaches and super-resolution imaging in fixed cells have revealed that there are ∼100-150 Xist molecules/cell that localise to around 60 preferred association sites (Xist hotspots) distributed across Xi^5–9^. Related to these findings, Xist hotspots are highly correlated with chromosome A1 sub-compartments that have a high 3D contact frequency with the Xist locus and with one another^7,10^, suggesting a role for 3D chromosome topology in directing Xist RNA localisation. Additionally, specific Xist RNA-binding proteins (RBPs) have inferred roles in Xist RNA localization. Key examples are the RBPs HNRNPU, proposed to tether Xist molecules to Xi chromatin^10,11^, HNRNPK implicated in retaining Xist RNA over Xi via formation of liquid-liquid phase separated (LLPS) droplets^12^, and SPEN, initially identified as a key silencing protein in XCI, but subsequently shown to have a genetically separable function in Xist RNA localisation^9^. Additionally, a number of Xist RBPs that bind to the Xist E-repeat sequence element, specifically CIZ1, MATR3, PTBP1, TDP43 and CELF-1, contribute to Xist RNA localisation during maintenance of XCI in differentiated cells^13,14^. This distinct pathway has also been suggested to rely on formation of an LLPS Xi compartment^14^.

The aforementioned advances have provided a framework for developing models to explain the unique behaviour of Xist RNA, but to date we have lacked the means to rigorously test these ideas. A key limitation is our inability to observe and quantify the dynamic behaviour of individual Xist RNA molecules either in the presence or absence of candidate localization factors. In this study we address this challenge by exploiting Nuc-BRITE, a novel approach for live-cell super-resolution imaging of nuclear RNAs at fast frame rates^15^, coupled with approaches to acutely deplete specific localisation factors. Using this strategy, we demonstrate that Xist molecules retain their chromatin association at all times throughout interphase, and that Xist spreading results from handover events linked to high frequency 3D-chromosome contacts. This contact-mediated transfer mechanism depends critically on the Xist RBPs HNRNPU and SPEN.

### Fast live-cell imaging demonstrates constrained behaviour of Xist RNA molecules

We developed Nuclear-Background Reduced RNA Imaging using Tamoxifen and ERT2 (Nuc-BRITE) to enable fast super-resolution (SR) imaging of RNA in the cell nucleus^15^. For this study we engineered an MS2v7 x24 array into the doxycycline inducible Xist allele of the previously described interspecific XX mouse ES cell (mESC) line iXist-ChrX^129^ ^16^. We then stably integrated a transgene encoding a pCAG driven Nuc-BRITE construct (tdMCP-tdStayGold-ERT2) (Fig 1a and b) which utilizes an ERT2 domain to reduce excess fusion protein in the cell nucleus. To visualize the Xist transcription site (TS) together with Xist RNA we further modified the Nuc-BRITE mESC line with a 100x LacO array^17^ located ∼ 3 kb downstream of Xist, and a transgene encoding LacI**-tdTomato integrated into the Rosa26 locus (Fig 1a and b). Finally, we integrated a pCAG driven Halo-PCNA construct to analyze the dynamic behaviour of Xist RNA in relation to DNA replication in S-phase (Fig 1a and b). At each genome engineering step we used allelic ChrRNA-seq analysis^16^ to confirm retention of X-linked gene-silencing capacity and levels of Xist RNA (Supplementary Fig 1a), and quantitative analysis of Xist RNA-FISH to verify normal Xist RNA localization patterns (Supplementary Fig 1b - e).

**Fig. 1.**
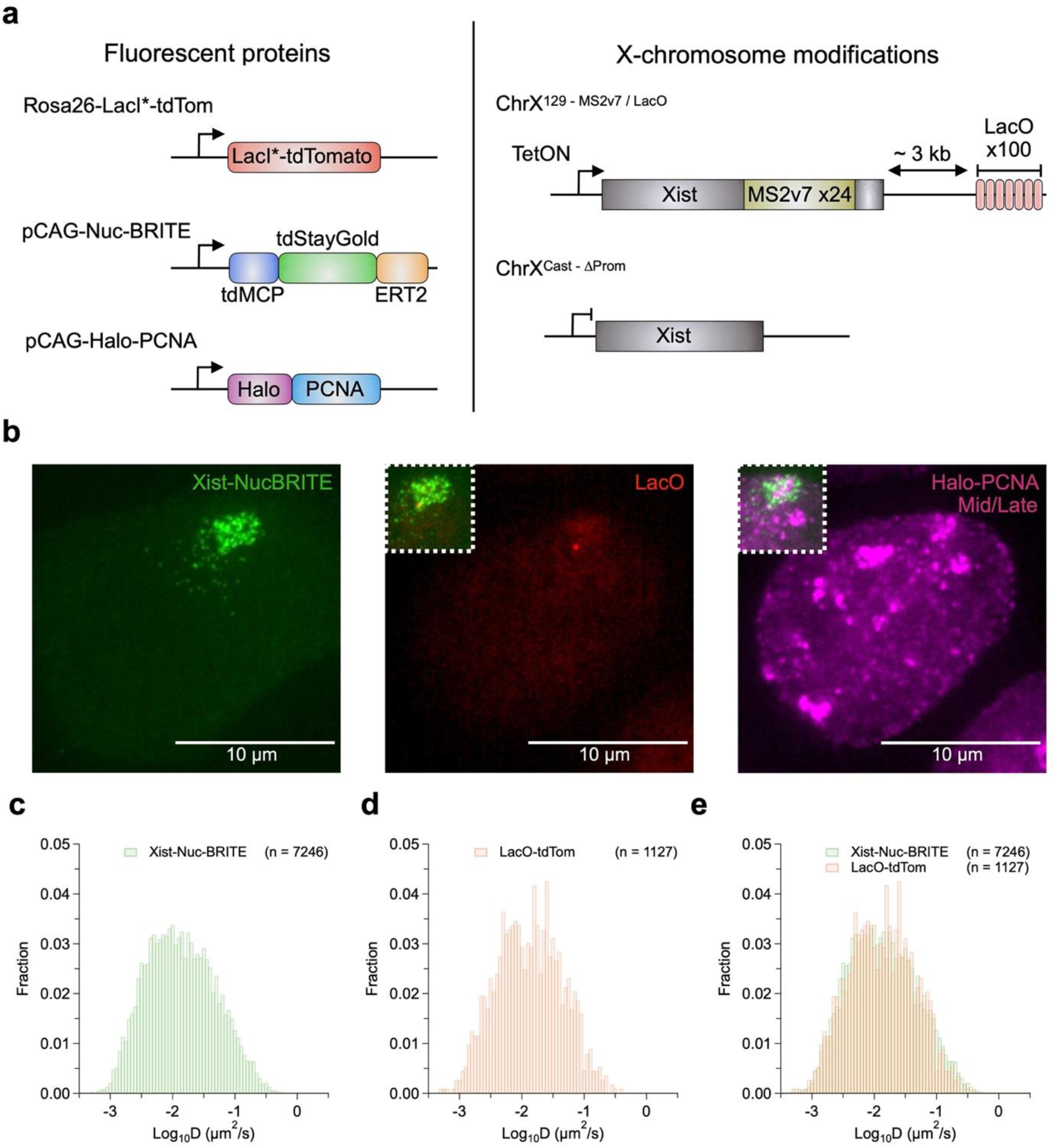
Constrained dynamic behaviour of Xist RNA determined using Nuc-BRITE. (a) Overview of genetic modifications in the Xist Nuc-BRITE cell line depicting fluorescent proteins and Xi modifications for MS2v7 and LacO arrays. (b) Max intensity projection of fixed Xist Nuc-BRITE cells using the VisiTech iSIM microscope showing Xist signal - green, LacO signal - red and Halo-PCNA - magenta from a cell in mid/late S-phase. Insets show overlays of LacO and Halo-PCNA signal in relation to Xist. (c) Log_10_ diffusion coefficient histograms for Xist Nuc-BRITE and (d) LacO-tdTom and (e) both overlaid. Number of tracks analysed were 7246 and 1127 taken from 319 and 443 cells respectively, from 3 independent replicates each.

The enhanced signal-to-noise ratio achieved with Nuc-BRITE enabled improved single particle tracking (SPT) analysis of Xist foci using fast frame rates (75 ms)^15^. In initial experiments we used SPT to measure diffusion coefficients (D) of individual Xist foci 24 h after induction with doxycycline. Log_10_D values (µm^2^/s) describe the dynamic behaviour of molecules, with values > −0.5 being indicative of unbound/diffusive behaviour and values < −0.5 indicating bound/constrained molecules. Strikingly, we observed that Xist molecules are for the most part constrained, with few if any mobile/diffusive molecules (Fig 1c, Supplementary Video 1). Moreover, the dynamic behaviour of Xist RNA was indistinguishable from that of the TS LacO array (Fig 1c - e), indicating that Xist RNA mobility reflects that of molecules tethered to the underlying chromatin.

### Dynamic behaviour of Xist RNA through the cell cycle

To determine if the constrained behaviour of Xist molecules observed in unsynchronized mESCs varies through the cell cycle we determined Xist log_10_D values at specific cell cycle stages, stalling cells in G2 with the Cdk1 inhibitor (RO-3306) and then analyzing Xist dynamics by SPT at G2 (RO-3306 present), and upon release, in mitosis and post-mitotic G1 stages. To analyze S-phase cells, we used the Halo-PCNA signal to classify early, mid and late S-phase cell populations (Fig. 2a). Log_10_D in G1 or S-phase cells were indistinguishable from those in unsynchronized cells, including for early-, mid- and late-S sub-populations (Fig 2b and c and Supplementary Fig 1, f - h). A small shift in log_10_D within the boundaries of chromatin associated movements was seen for G2 cells (Fig 2d). A more significant change was observed at mitosis with the appearance of a large population of unbound or diffusive Xist molecules with log_10_D values > −0.5 (Fig 2e). This latter observation is consistent with prior studies demonstrating dissociation of Xist from the Xi chromosome during mitosis^18–21^. Importantly, capturing diffusive behaviour of Xist RNA at mitosis underscores that the highly constrained chromatin-like dynamics seen at other cell cycle stages is biological rather than a technical limitation in tracking fast diffusing molecules.

**Fig. 2.**
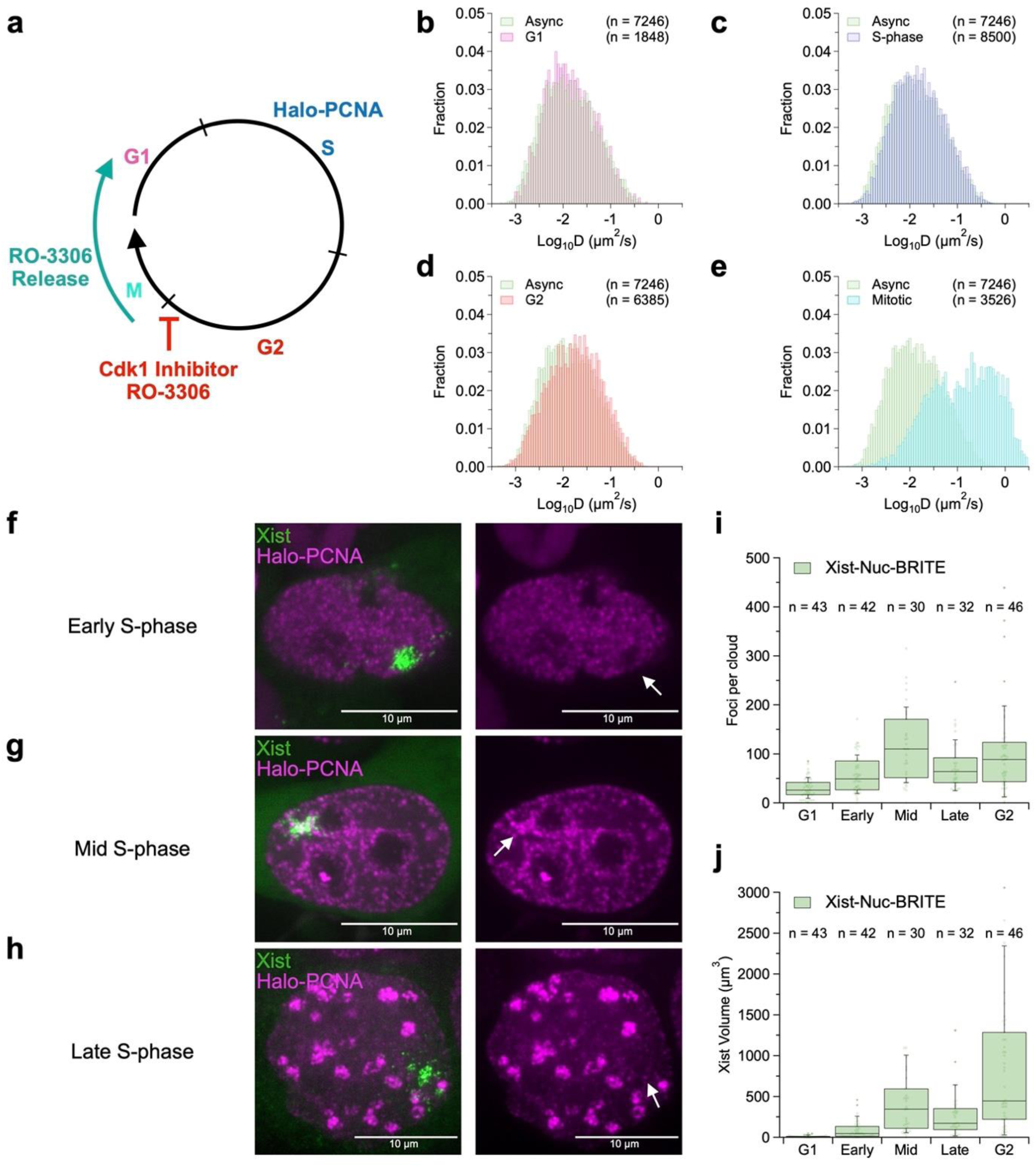
Dynamic behaviour of Xist RNA through the cell-cycle. (a) Diagram of cell-cycle inhibition (16h - RO-3306) and release allowing the separation of cell-cycle phases into G1, S, G2 and Mitosis. (b - e) Log_10_ diffusion coefficient histograms for (b) G1 - magenta, (c) S-phase - blue, (d) G2 - red and (e) Mitosis - cyan, all compared to asynchronous cells acquired from all stages within interphase. (f - h) Representative Z-sections of Halo-PCNA signal in Xist Nuc-BRITE cells (Xi domain marked with arrows) in (f) early S-phase, (g) mid S-phase and (h) late S-phase. (i and j) Quantification of number of Xist foci per cell (i) and volume of Xist foci per cell (j) from G1 through to G2 phases, data taken from fixed Xist Nuc-BRITE cells with n representing number of cells quantified.

Interestingly, although Xist showed no difference in diffusive behaviour during S-phase, it was evident that the distribution of Xist molecules and the volume that they occupy does change. To investigate this observation, we quantified the number Xist foci and the volume of Xist territories in fixed cells at different stages of the cell-cycle (Fig 2f - j). Numbers of Xist foci and territory volumes were relatively low in G1, consistent with their reestablishment in post-mitotic cells. During early S-phase Xist territories were larger but relatively compact. Halo-PCNA signal was absent from the Xi, indicating replication has not commenced. During mid S-phase, Xist territories were more dispersed with associated increases in the number of individual foci and the territory volume. Additionally, Halo-PCNA could now be seen within the Xist territory, indicating ongoing Xi replication (Xi replication in mid-S-phase is in line with prior work^22^). Given that Xist mobility does not increase in mid-S-phase, we infer that increased dispersal is a consequence of a disruption of higher order chromosome organization during synchronous Xi DNA replication. In late S-phase, the number of Xist foci and the volume they occupy is reduced, likely reflecting post-replicative reinstatement of higher order chromosome structure.

### A contact-mediated transfer model for Xist RNA spread

Quantitative analysis of Xist molecules using SR imaging in fixed cells indicates that Xist multimerizes at sites along the X chromosome, with individual Xist foci comprising on average 2 Xist molecules^8,9^. Consistent with these observations, we noted variability in the intensity of individual Xist foci resolved using Nuc-BRITE imaging. To quantify this parameter, we co-transfected the Xist Nuc-BRITE mESCs with a plasmid that transcribes MS2v7 x24 alone and then used the intensity of fluorescent cytoplasmic particles to define a value for single RNA molecules (Fig 3a, Supplementary Video 2). We then used this value to calibrate the intensity of Xist Nuc-BRITE foci at interphase. As shown in Figure 3a we observed a broad distribution of intensities corresponding to 1-6 molecules with a mean value of 2, consistent with previous observations^8,9^.

**Fig. 3.**
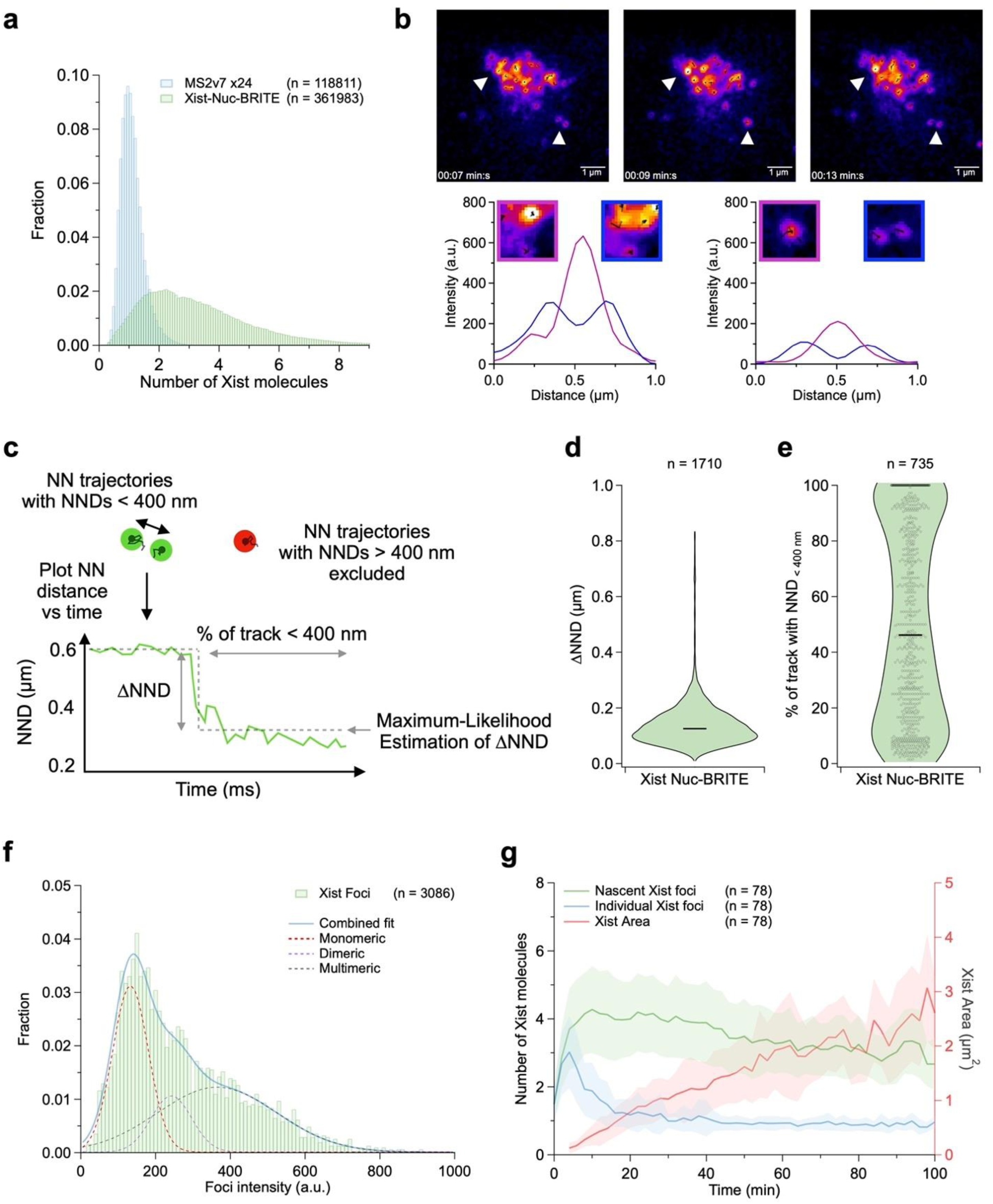
Quantification of Xist foci accumulation and interactions. (a) Number of Xist molecules per foci, normalized from MS2v7 x24 intensity distribution - blue compared to Xist Nuc-BRITE multimeric accumulation - green. N values depicted as number of foci taken from 50 and 120 cells for MS2v7×24 and Xist Nuc-BRITE respectively. (b) Representative snapshots of movie (Supplementary Video 3) from fast frame rate imaging with 24h of dox induction showing Xist-Xist interactions over time. Intensity profiles depict higher intensity fused foci - magenta and lower intensity split foci - blue for both large and small accumulations of Xist. (c) Diagram showing the principles behind selection of nearest neighbor trajectories, plotting of NNDs over time and maximum likelihood fitting of NND traces to calculate ΔNND and durations of interactions. Violin plots of ΔNND values (d) and percentage of trajectories < 400 nm (e) for wild-type Xist Nuc-BRITE cells. N values depicted as number of ΔNND values and number of trajectories analyzed for d and e respectively from a total of 319 cells from 3 independent replicates. (f) Histogram of Xist Nuc-BRITE foci intensities from 2h time course following Xist induction (n = 3086 foci from 78 cells) fit with a triple gaussian fit to represent different Xist intensity populations and single gaussian fits to depict monomeric, dimeric and multimeric intensities. (g) Average number of molecules for nascent - green and individual Xist foci - blue as well as Xist territory area - red plotted over time with the standard deviation plotted as the shaded area around the mean. Data taken from 78 cells.

In prior work using RNA-SPLIT we reported that newly synthesized Xist molecules predominantly co-localize to sites occupied by previously synthesized molecules, termed coupling^9^, indicating that Xist does not translocate as a multimer, but rather multimerizes at preferred association sites along the X chromosome. Given that Xist molecules are non-diffusive and apparently chromatin bound throughout interphase, and taking into account work showing that preferred association sites correspond to chromosomal locations that have a high contact frequency with the Xist locus and with one another^7,10^, we invoke that spread of Xist molecules along the chromosome must occur through contact-mediated transfer in which chromosome interactions provide an opportunity for bound Xist molecules to relocate to a neighboring site. In support of this conclusion, fast imaging with Nuc-BRITE following Xist induction for 24h reveals that neighboring Xist foci undergo frequent fusion and splitting events with associated intensity changes (Fig 3b, Supplementary Video 3). To quantify this behaviour, we analyzed changes in nearest neighbor distances (NND) for paired Xist foci over time leading up to fusion and splitting events (Fig 3c, Supplementary Note 1). First, we quantified the distance that Xist pairs move towards or away from one another (ΔNND), which was observed to be ∼150 nm on average (Fig 3d). Second, we measured the duration of interactions within a given trajectory by quantifying the percentage of the track where pairs were less than 400nm apart. Thus, we observed a distribution of tracks with the most stable interactions showing ∼100% of the trajectory < 400 nm apart and more dynamic interactions with foci spending ∼10% of the trajectory < 400 nm apart (Fig 3e). These parameters provide a basis to compare dynamic interactions of Xist foci under different conditions or following perturbation of specific factors (see below).

To further investigate how Xist RNA spreads we analyzed the expansion of Xist RNA territories during an initial period of 2h following doxycycline induction, exploiting LacO/LacI labelling to pinpoint the Xist TS. By imaging with a relatively slow frame rate (2 min) we could observe an accumulation of nascent Xist molecules, followed by spreading, evident as dispersed foci with lower intensity values (Supplementary Fig 2a and Supplementary Video 4). Tracking Xist foci with the highest and lowest intensities allowed recording of nascent Xist accumulations and isolated Xist foci over time. Plotting these intensity trajectories and the Xist territory area shows the relationship between transcriptional activity and the commencement of Xist spreading (Supplementary Fig 2b). As noted for Xist intensity distributions after 24h of induction (Fig 3a), mean intensity values over the 2h following dox induction similarly indicate most foci comprising either single or two Xist molecules (Fig. 3f). The larger TS foci on the other hand quantified as ∼3–8 molecules of Xist RNA. Tracking both TS and the lowest intensity foci detected over time shows, on average, TS accumulations persist for ∼ 5-10 mins after which Xist starts to spread (Fig. 3g). Further bursts of Xist transcription led to additional large accumulations of Xist RNA and subsequent spreading over a similar ∼10 mins timescale, with an overall spreading rate of ∼2 µm^2^/h, correlating well with the spreading rate estimated using fixed cell analysis^9^. Using fast imaging during the initial 2h period after Xist induction we again observed frequent fusion and splitting of Xist foci (Supplementary Fig 2c and Supplementary Video 5). We infer that these events represent handover of Xist molecules between chromatin contact sites, as argued above for analysis of Xist interactions that lead to fusion and splitting of foci 24h after Xist induction (Fig 3b).

### SPEN and HNRNPU constrain Xist RNA at interphase

Prior studies identified the Xist RBPs SPEN, HNRNPK and HNRNPU as having roles in Xist RNA localization during establishment of XCI ^9,11,12,23^. To investigate how these factors contribute to Xist RNA dynamic behaviour we engineered Xist Nuc-BRITE mESCs to deplete each factor/pathway: For SPEN, which is not required for mESC viability^16,24,25^, we used CRISPR/Cas9 mediated mutagenesis to generate a line with null alleles (SPEN^KO^) and a second mutant with an in-frame deletion spanning sequences encoding SPEN RRM domains 2-4 required for binding to the Xist A-repeat region (SPEN^ΔRRM^)^24,26^ (Fig 4a, Supplementary Fig 3a). HNRNPK binds to the Polycomb interaction domain (PID) within the B/C tandem repeat region, and we therefore made use of a deletion of this region (Xist^ΔPID^)^16,27^ in Xist Nuc-BRITE mESCs to investigate loss of HNRNPK recruitment on Xist RNA localization (Fig 4a, Supplementary Fig 3b). Additionally, we engineered an FKBP12^F36V^ tag on the N-terminus of HNRNPK (dT-hnK) to enable acute depletion following addition of dTAG-13 to the culture medium (Fig 4a, Supplementary Fig 3c). Finally, an FKBP12^F36V^ tag was engineered onto the C-terminus of HNRNPU (hnU-dT) to enable acute depletion following addition of dTAG13 (Fig 4a, Supplementary Fig 3d).

**Fig. 4.**
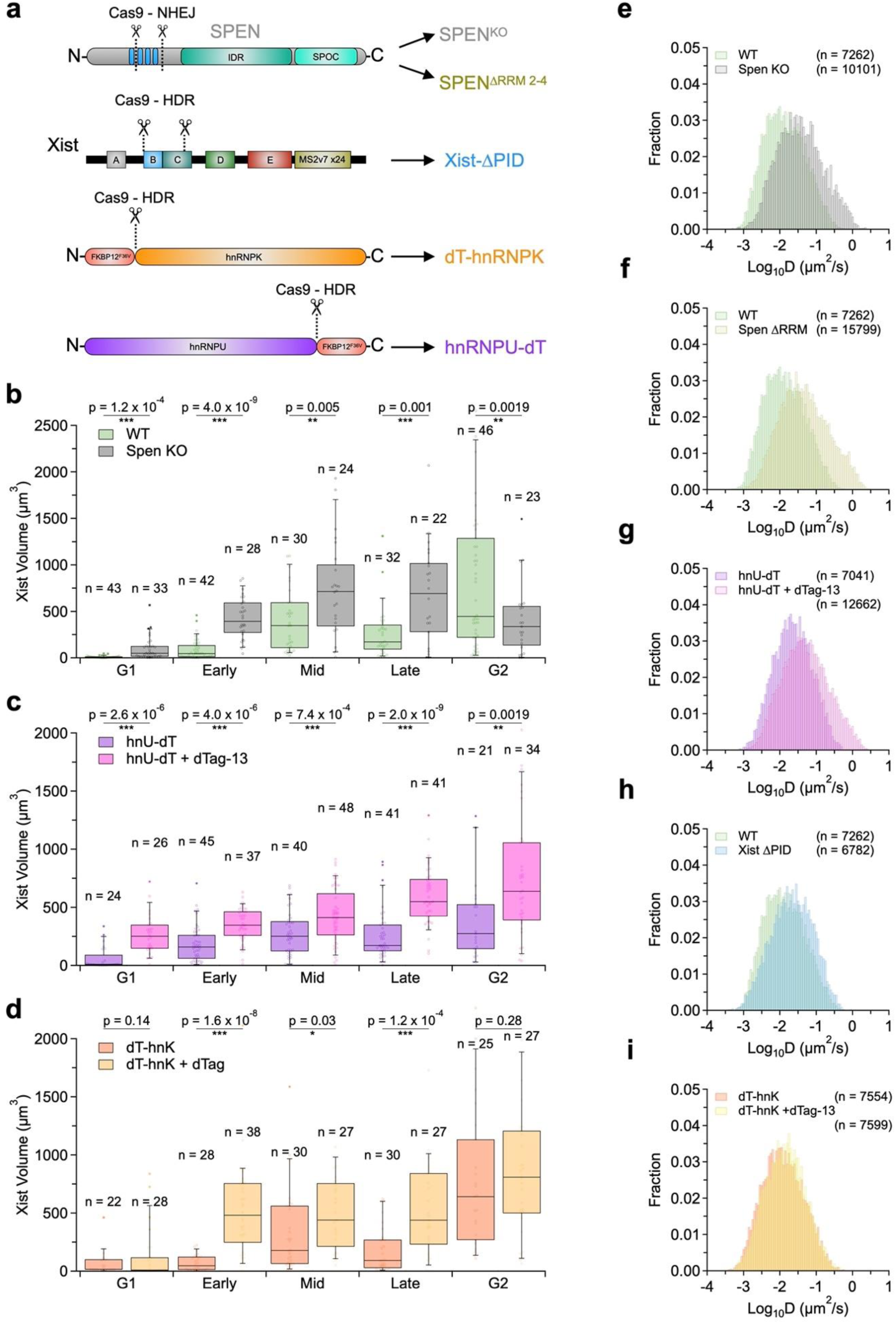
Contribution of Xist localization factors to Xist RNA dynamic behaviour. (a) Diagram of adaptations to Xist Nuc-BRITE cell lines for each mutant cell line generated with annotated CRISPR/Cas9 strategies for non-homologous end joining (NHEJ) and homologous directed repair (HDR). (b - d) Quantification of Xist volume from fixed cells based on Halo-PCNA signal and separated into cell-cycle phases G1 through to G2. Xist Nuc-BRITE data was compared to SPEN ^KO^ (b), hnU-dT compared to hnU-dT + dTAG-13 and (c) and dT-hnK compared to dT-hnK + dTAG-13 (d). Numbers of cells analyzed per cell-cycle stage displayed as n. (e - i) Log_10_ diffusion coefficient histograms for each mutant cell line either compared to wild-type Xist Nuc-BRITE (WT) or no dTAG-13 control cells in dTAG lines. Number of trajectories analyzed presented as n from 319, 361, 363, 213, 301, 324, 258 and 212 cells from WT, SPEN ^KO^ (e), SPEN ^ΔRRM^ (f), Xist-ΔPID (g), dT-hnK, dT-hnK + dTAG-13 (h), hnU-dT and hnU-dT + dTAG-13 (i) lines respectively using 2 to 3 replicates per cell line.

All modified lines were validated using allelic ChrRNA-seq, confirming previously reported effects on X-linked gene silencing and Xist RNA levels^9,10,16^ (Supplementary Fig 4a and b). Aberrant Xist localization patterns were confirmed by examination of Xist Nuc-BRITE signal after 24h Xist induction in unsynchronized cells. Each mutation gave the expected Xist localization phenotype 24 h after Xist induction: SPEN^KO^ and SPEN^ΔRRM^ resulted in smaller Xist clouds with widely dispersed Xist foci. Xist-ΔPID and dT-hnK + dTAG-13 generated larger more dispersed Xist clouds, and hnU-dT + dTAG-13 led to significant dispersal of Xist molecules through the nucleoplasm (Supplementary Fig 4c, Supplementary Video 6). For selected mutants we quantified Xist territory volumes in fixed cells at different cell cycle stages using RO-3306 synchronization as described above (Fig 4b - d). Thus, SPEN^KO^ cells up to G2, have increased territory volumes, in line with Xist RNA dispersal reported previously^16^. Similarly, HNRNPU depletion results in larger Xist territories at all cell cycle stages, as reported previously^11^. HNRNPK depletion resulted in expanded Xist territories from S-phase onwards, again consistent with prior studies^23^.

We next determined the role of the different localization factors in Xist dynamic behaviour using SPT analysis after 24h Xist induction with and without depletion of individual factors/pathways (Fig 4e - i, Supplementary Video 6). Thus, we plotted log_10_D histograms and compared to a control of either wild-type Nuc-BRITE cells (WT) in constitutive mutants or no dTAG-13 treatment in the HNRNPK and HNRNPU degron lines. Both SPEN^KO^ and SPEN^ΔRRM^ mutants show a significant proportion of diffusive molecules with a clear increase in D, indicative of dissociation of Xist molecules from chromatin (Fig. 4e and f). This population of molecules have log_10_D values that are in the same range as diffusive Xist particles observed in mitotic cells (Fig 2e). Similarly, in hnU-dT cells treated with dTAG-13, a significant population of Xist molecules are diffusive, indicating loss of chromatin association (Fig. 4i).

In contrast to perturbations of SPEN and HNRNPU, abrogation of HNRNPK function (Xist^ΔPID^ and dT-hnK cells treated with dTAG-13) had relatively subtle effects on Xist dynamic behaviour. Thus, in Xist^ΔPID^ cells there is a small shift in D toward the right, with a reduction of the highly immobile fraction of particles with values less than −2 (Fig 4g). Log_10_D values for the majority of molecules remain below −0.5, indicating Xist molecules are less restricted in their motion, but still display chromatin associated dynamics. A similar, but less prominent shift is seen in the degron HNRNPK cells treated with dTAG-13, again consistent with a continued association with chromatin (Fig 4h). Accordingly, we conclude that the apparent dispersal of Xist RNA following HNRNPK depletion is not linked to increased diffusive behaviour. Alternative explanations include elevated levels of Xist RNA seen in both Xist^ΔPID^ and dT-hnK cells treated with dTAG-13 (Supplementary Fig 4b). Additionally, we find evidence for advanced Xi DNA replication timing (Supplementary fig 5a and b, Supplementary Note 2) and a decrease in Xi chromatin compaction, likely due to abrogation of polycomb recruitment (Supplementary fig 5c - f, Supplementary Note 2), both of which could account for the modified appearance of Xist RNA territories.

### Distinct Roles for SPEN and HNRNPU in Xist RNA localization behaviour

We went on to investigate how perturbing Xist localization factors affects contact-mediated transfer, initially analyzing the dynamic interactions of Xist foci inferred to represent hand-over events transferring Xist molecules between neighboring sites. Fusion and splitting of Xist foci could be observed for all mutant backgrounds (Fig 5a - d), but quantitative analysis of Xist interactions leading to these events demonstrated clear differences (Fig 5e and f). Thus, we observed an increase in median ΔNND values from ∼ 150 nm up to ∼ 200 nm for SPEN^KO^, SPEN^ΔRRM^ and hnU-dT + dTAG-13 compared to respective controls. A smaller and less significant increase is seen for the Xist-ΔPID cell line, whilst dT-hnK+ dTAG13 cells showed no significant change. In relation to the duration of interactions of neighboring Xist foci, SPEN and hnU-dT + dTAG-13 treatment resulted in a marked decrease in the proportion of time that foci were interacting over each trajectory. Conversely, both Xist^ΔPID^ and dT-hnK+ dTAG13 cells show a small increase in the duration of interaction of neighboring foci.

**Fig. 5.**
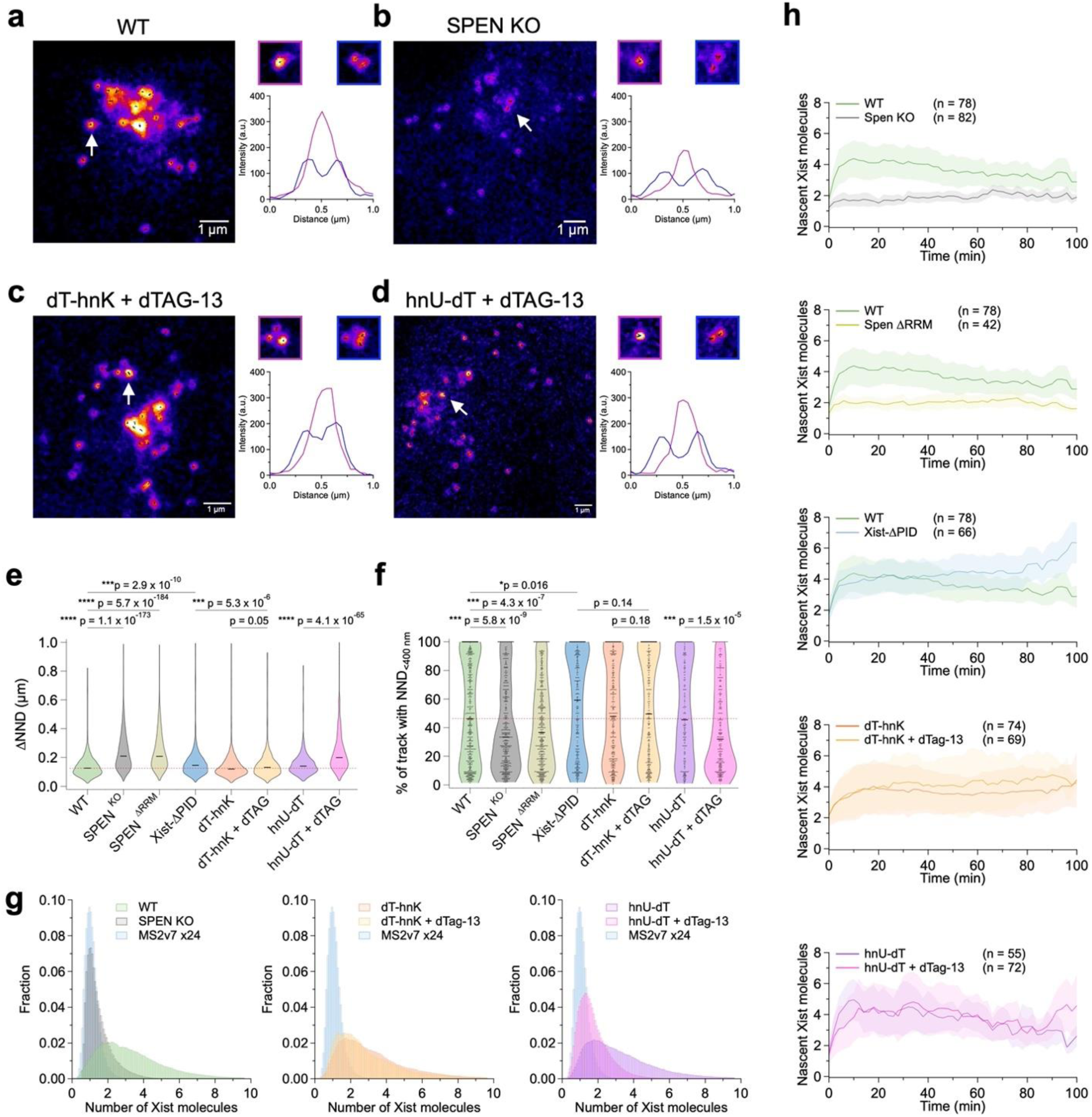
Separable functions for SPEN and HNRNPU in Xist RNA spread. (a - d) Representative images and intensity profiles of Xist accumulation and interacting foci taken from fast frame rate live-cell imaging for Xist Nuc-BRITE (a), SPEN ^KO^ (b), dT-hnK + dTAG-13 (c) and hnU-dT + dTAG-13 (d). (e) Violin plots of ΔNND values across all cell lines with the number of ΔNND values detected in each sample being; 1710 (WT), 2972 (SPEN ^KO^), 4148 (SPEN ^ΔRRM^), 893 (Xist-ΔPID), 1315 (dT-hnK), 1154 (dT-hnK + dTAG-13), 798 (hnU-dT ) and 2644 (hnU-dT + dTAG-13 ). Statistics shown from single tailed t-tests. (f) Violin plots of percentage duration of paired tracks with NND values less than 400 nm across all cell lines with the number of paired trajectories quantified being; 735 (WT), 644 (SPEN ^KO^), 678 (SPEN ^ΔRRM^), 418 (Xist-ΔPID), 509 (dT-hnK), 440 (dT-hnK + dTAG-13), 265 (hnU-dT) and 400 (hnU-dT + dTAG-13) and statistics used single tailed t-tests. Data for e and f was taken from 319 (WT), 361 (SPEN ^KO^), 363 (SPEN ^ΔRRM^), 213 (Xist-ΔPID), 301 (dT-hnK), 324 (dT-hnK + dTAG-13), 258 (hnU-dT) and 212 (hnU-dT + dTAG-13) cells using 2 to 3 replicates per cell line. (g) Number of molecules per foci normalized to monomeric MS2v7 x24 intensity distribution – blue. Numbers of foci analyzed were 118811 (MS2v7 x24 - blue), 361983 (WT - green), 236261 (SPEN ^KO^ - grey), 318522 (dT-hnK - orange), 384319 (dT-hnK + dTAG-13 - yellow), 473647 (hnU-dT - purple) and 415470 (hnU-dT + dTAG-13 - pink taken from 50, 120, 181, 113, 151, 149 and 107 cells respectively. (h) Average number of nascent Xist molecules over time following dox induction for each mutant cell line with comparison to wild-type cells or no dTAG-13 treatment where appropriate. Number of cells presented as n and standard deviation depicted as shaded area around the mean.

We also quantified focal Xist intensities in representative mutant backgrounds, providing a measure of Xist multimerization at preferred association sites (Fig 5g). dT-hnK + dTAG-13 cells show a broad distribution of focal intensities similar to wild-type cells (ranging from 1-8 Xist RNA molecules), indicating little or no change in Xist multimerization. In contrast, for SPEN^KO^ cells we observed a striking shift with most foci having intensities equivalent to monomeric or dimeric Xist molecules. hnU-dT + dTAG-13 cells also showed reduced focal accumulation, although not to the same degree as in SPEN^KO^ cells.

The analysis of focal Xist intensities provides an indication that SPEN and HNRNPU may have distinct functions in regulating Xist RNA spread. To further investigate this, we determined how the aforementioned perturbations affect Xist RNA spread at early timepoints following doxycycline induction. Data acquired from imaging Xist in the first 2 hours post-induction shows that only SPEN^KO^ and SPEN^ΔRRM^ affect the large accumulations of nascent Xist molecules seen in wild-type cells (Fig. 5h). Specifically, for both SPEN mutations the intensity of nascent Xist accumulations at the TS is maintained at a level equivalent to 1-2 Xist molecules compared to foci approximating to 4-6 Xist molecules as seen in wild-type cells and following depletion of other localization factors. Thus, taken together our observations demonstrate key roles for both HNRNPU and SPEN in Xist RNA spread. The former facilitating the tethering of Xist to chromatin and the latter facilitating focal accumulation of Xist RNA at the TS and preferred association sites.

## Discussion

The application of Nuc-BRITE and fast live-cell imaging enabled measurement of the dynamic behaviour of Xist RNA, allowing us to show definitively that the majority of Xist molecules are chromatin/chromosome bound throughout interphase. This conclusion is supported by the observation that Xist RNA is measurably unbound in mitotic cells, when Xist RNA is known to dissociate from Xi, and as a result of mutations in the Xist RBPs SPEN and HNRNPU. Our findings are in accordance with biochemical experiments that show Xist RNA is in large part bound up in the nuclear matrix or insoluble/chromatin bound fraction^18^. The absence of a diffusive population of Xist molecules during interphase leads us to the conclusion that Xist translocation occurs via a handover mechanism when distant sites make 3D contacts with the Xist locus or with one another, referred to herein as contact-mediated transfer (Fig 6a). This proposal integrates prior work showing that high frequency intrachromosomal contact sites for the Xist locus, determined using HiC analysis, correlate with preferred sites for Xist RNA localization^7^. In recent work we report that these sites correlate with A1 sub-compartments on the X chromosome which interact both with the Xist locus and with one another^10^. Whether it is the frequency or duration of such contacts that enables Xist translocation events remains to be determined, and this will be an interesting issue to investigate in future work. Contact-mediated transfer also provides an explanation for our previous finding that newly synthesized Xist molecules predominantly co-localise with pre-synthesized molecules^9^.

**Fig. 6.**
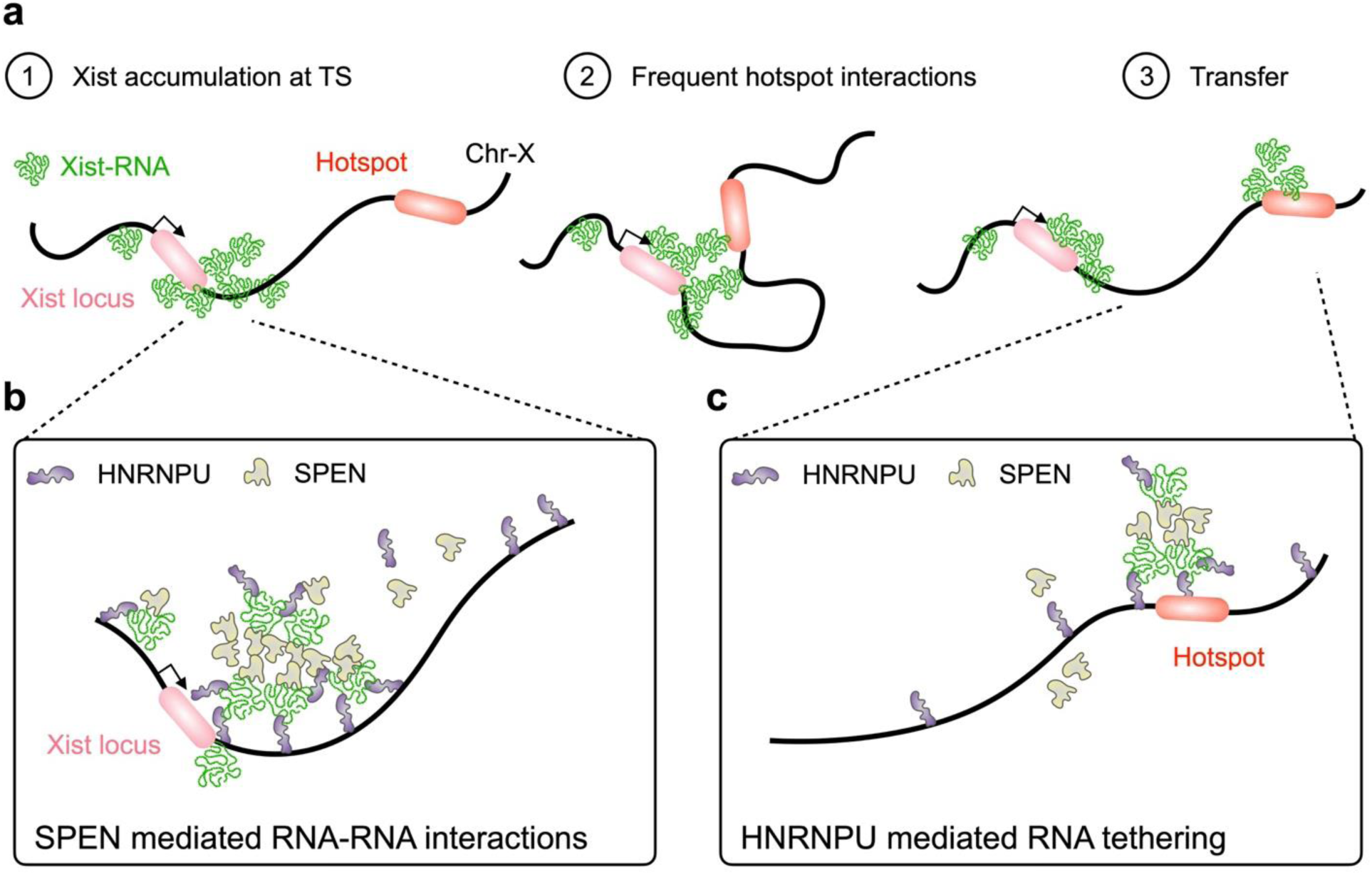
Model of Xist RNA spreading via contact-mediated transfer. (a) Diagram of the initial stages of Xist RNA spreading shown as Xist RNA (green) accumulation at the Xist locus (pink), followed by frequent interactions with a neighboring Xist hotspot (red), leading to the transfer of Xist to new sites on the Xi. Diagram showing proposed Xist RNA tethering and RNA-RNA interactions facilitated by HNRNPU and SPEN respectively at the Xist transcription site (TS) (b) and Xist hotspot (c).

It was recently proposed that HNRNPK has a key function in Xist RNA spread via an LLPS mechanism^12^. However, we found that HNRNPK depletion (and similarly deletion of the Xist PID region) results in only minor changes in the dynamic behaviour of Xist RNA, with the majority of molecules exhibiting constrained mobility indicating continued chromatin association. We did observe enlargement and relative dispersal of Xist RNA territories, consistent with prior reports^12,23^. We note that dispersal of Xist territories also occurs upon depletion of the paralogous Polycomb proteins PCGF3 and PCGF5, which interact with HNRNPK to initiate the Polycomb pathway in XCI^28^, suggesting that Xist dispersal occurs downstream of HNRNPK. Similarly, the deficit in Xi silencing following deletion of the PID region and PCGF3/5 is equivalent, again arguing against a function for HNRNPK other than recruiting Polycomb to facilitate X-linked gene silencing and Xi chromosome compaction^27^. With these considerations in mind, we propose reduced compaction of Xi chromatin as an alternative explanation for Xist dispersal in these scenarios. The observed advance in Xi replication timing and/or increased levels of Xist RNA may also be contributory factors.

Depletion of HNRNPU resulted in a clear shift in the dynamic behaviour of Xist RNA with the appearance of a diffusive population during interphase. We attribute this to HNRNPU function in tethering Xist molecules to the chromosome through simultaneous interaction with RNA targets and DNA/chromatin via C-terminal RRM and N-terminal SAP domains respectively^11^. Localization of Xist molecules to preferred association sites and to some degree Xi silencing are retained following HNRNPU depletion^10^, and accordingly we propose that translocation via contact-mediated transfer does occur, but that the duration of chromosome binding is reduced, enabling Xist molecules to detach and diffuse to non-Xi sites (Fig 6b and c). A fraction of Xist molecules retain constrained dynamic behaviour following HNRNPU depletion, indicative of chromatin association. The basis for this finding is presently unknown but may reflect the contribution of other chromosome tethering factors, for example CIZ1 and associated proteins, functioning synergistically with HNRNPU.

SPEN loss of function mutations, like HNRNPU depletion, result in the appearance of a diffusive population of Xist molecules. This finding is in accordance with our prior work demonstrating aberrant Xist localization as a result of SPEN loss of function^9^. Here we invoke a different underlying mechanism to that suggested for HNRNPU depletion. Thus, the initial accumulation of Xist molecules at the transcription start site, and the formation of multimeric Xist foci, are both strongly abrogated, suggesting a reduction in Xist-Xist vs Xist-chromatin interactions. This conclusion is reinforced by our prior observation that Xist coupling, as measured using RNA-SPLIT, is largely absent with SPEN loss of function^9^. Of note, mutation of the SPEN SPOC domain required for interaction with NCoR-HDAC3 and downstream Xi hypoacetylation and X-linked gene silencing does not affect Xist localization, implicating other domains in the SPEN protein^9^. An interesting candidate is SPEN’s extensive intrinsically disordered regions which have been suggested to mediate weak homotypic SPEN-SPEN interactions^8^. Accordingly, we speculate that SPEN-SPEN interactions facilitate Xist-Xist interactions that in turn contribute to the formation of Xist accumulations at the transcription site and at preferred association sites, thus facilitating Xist translocation via contact-mediated transfer (Fig 6b and c).

To summarize, the application of fast live-cell imaging of Xist RNA using Nuc-BRITE has provided key insights into the dynamic behaviour of Xist RNA during XCI. Together with observations from related studies, this has allowed us for the first time to develop a rational and coherent model to explain *cis-*limited spread of Xist RNA across the chromosome from which it is transcribed.

## Methods

### Cell Culture

Mouse embryonic stem cells (mESCs) were cultured at 37°C and 5% CO2 in ESC media: High Glucose Dulbecco’s Modified Eagle Medium no pyruvate (Life Technologies) supplemented with 10% Foetal Bovine Serum (Sigma), 0.1mM non-essential amino acids, 2mM L-glutamine, 50μM β-mercaptoethanol, 100 U/ml Penicillin/100 μg/ml Streptomycin (Life Technologies) and 1000U/mL Leukemia Inhibitory Factor (LIF) (made in-house). mESCs were grown on gelatinised plates, on mitotically arrested mouse SNLP feeder cells (STO mouse fibroblasts expressing Neomycin, Puromycin resistance and LIF genes). Stocks of feeders were frozen down from mitotically arrested SNLPs cultured in mitomycin-C (Sigma aldrich) at 1µg/ml for 2h. For imaging experiments, mESCs were grown in gelatinised ibidi 8-well chamber slides in the absence of feeders and imaged in live-cell imaging media: FluoroBrite DMEM supplemented with 20 mM HEPES, 10% Foetal Bovine Serum, 0.1mM non-essential amino acids, 2mM L-glutamine, 50μM β-mercaptoethanol, 100 U/ml Penicillin/100 μg/ml Streptomycin (Life Technologies) and 1000U/mL Leukemia Inhibitory Factor (LIF). Treatment of mESCs with doxycycline (Sigma) or dTAG-13 (Tocris Bioscience) used 1 µg/ml and 100 nM concentrations respectively. For mESC lines with Xist localization factor degron tags (see below), cells were cultured in ESC media and incubated in 100nM dTAG-13 two hours prior to dox induction and sustained for a further 24 h throughout XCI.

### Generation of Nuc-BRITE cell line

The previously described female 129/Cast interspecific iXist-ChrX^129^ mESC cell line^16^ was used as the parental cell line for this study. Multiple rounds of CRISPR/Cas9 engineering were used to generate the Nuc-BRITE cell line. CRISPR sgRNAs and screening primers can be found in Supplementary Table 1. Firstly, LacO-tdTom cells were generated by CRISPR/Cas9 homology directed recombination (HDR) via targeting a 100x LacO array 3kb downstream of Xist using a 129 strain specific sgRNA (doxycycline inducible Xist expression is on the 129 strain X chromosome). In the same transfection, Rosa26 was targeted with a Rosa26 specific sgRNA and Rosa26-LacI**-tdTomato plasmid^17^ (LacO plasmid was a gift from Luca Giorgetti, FMI, Switzerland). Cells were transfected in 6-well plates with lipofectamine 3000 following the manufacturers protocol with a total of 1.5µg of sgRNA plasmids and a 6:1 molar ratio of HDR plasmids. Following 24h of plasmid expression, cells were split and seeded into three densities, 1:2, 1:3 and 1:6 on 10cm dishes. After a further 24h incubation, puromycin was used for selection at a concentration of 3.5µg/ml for a further 48h. Following puromycin selection cells were recovered for ∼ 10 – 14 days at which time single colonies were picked and seeded into 96-well plates. Positive Rosa26-LacI-tdTom cell lines were initially screened by PCR and then using Xist RNA-FISH to confirm Xi-specific LacO-tdTom foci. Selected clones were tested for puromycin sensitivity.

The LacO-tdTom cell line was transfected as above to introduce an MS2v7 x24 array^29^ into Exon 7 of Xist at a position where introduction of a stem-loop array was previously shown not to alter Xist function and behaviour^9^. As above, clones were screened by PCR and RNA-FISH to confirm Xi-specific MS2 incorporation and an XX karyotype. LacO-Xist-MS2v7 cells were transfected with 1µg of the pCAG-tdMCP-tdStayGold-ERT2 or pCAG-NLS-tdMCP-tdStayGold plasmids and cells were selected with Neomycin (300µg/ml) for 14 days until single colonies formed. Individual clones were screened by PCR for transgene insertion and imaged live to detect efficient transgene expression and Xist-MS2 specific signal. Finally, Xist-Nuc-BRITE cells were transfected with a pCAG-Halo-PCNA plasmid, seeded at a low cell density and selected for with Neomycin initially at high concentration and reducing over 14 days of selection (600 – 300µg/ml). Single colonies were screened by PCR for transgene insertion and imaged live to detect Halo-PCNA signal using Halo-JFX650 labelling following the manufacturers protocol. Normal commencement of XCI was confirmed at each step using ChrRNA-Seq and allelic X-chromosome expression analysis after 24h dox induction, as described previously^16^.

### Generation of SPEN KO, SPEN ΔRRM, Xist-ΔPID, dT-hnK and hnU-dT cell lines

CRISPR/Cas9 KO and HDR strategies were used to modify Xist-Nuc-BRITE Halo-PCNA cells to introduce specific mutations or tags. CRISPR sgRNAs and screening primers can be found in Supplementary Table 1. SPEN KO and SPEN ΔRRM cells were generated by transfecting with Spen targeting guides sgRNA_3 and sgRNA_5 as previously described^16,24^. This transfection generated multiple different genotype clones via NHEJ which were screened by PCR for loss of Exon 3 downstream of sgRNA_3. Clones were then subject to Sanger sequencing and ChrRNA-Seq to confirm that the stable Spen mRNA products were either out-of-frame or in frame for SPEN KO and SPEN ΔRRM clones respectively. Furthermore, the phenotypic changes in Xist RNA spreading were confirmed using live-cell imaging. Xist-ΔPID clones were produced using CRISPR/Cas9 HDR with a single sgRNA targeting the Xist-B repeat and an Xist-ΔPID homology vector to repair the break to accurately introduce the full ΔPID deletion as described previously^16^. Accurate PID deletions were detected using PCR and confirmed with sanger sequencing and ChrRNA-Seq analysis. Additionally, the effect on Xist RNA spreading was confirmed with live-cell imaging and the loss of H2AK119ub (uH2A – Cell Signaling Technologies - #8240) deposition was confirmed with immunofluorescence. To acutely deplete the essential RBPs HNRNPK and HNRNPU we used the dTAG degron system^30^, engineering homozygous FKBP12F36V knock-in clones for HNRNPK and HNRNPU at the N-terminus and C-terminus respectively to generate dT-hnK and hnU-dT cell lines. Positive clones were screened via western blot using HNRNPK (CST - #4675) and HNRNPU (in house) antibodies to visualise homozygously tagged proteins. Additionally, positive clones were subject to 2h and 26h dTAG-13 (100nM) treatments to confirm loss of the tagged protein via western blot. The effect of HNRNPK and HNRNPU depletion on XCI was also assessed using ChrRNA-Seq and allelic analysis of gene expression.

### RNA-FISH and immunofluorescence

mESCs were grown on gelatin-coated No. 1.5H precision coverslips (170 ± 5 µm; Marienfeld Superior) in 6-well plates. Xist was induced with 1 µg/ml doxycycline for 24h. Coverslips were washed with PBS and fixed with 3.7% formaldehyde in PBS for 10 min at room temperature. Cells were rinsed three times with PBST.5 (PBS + Tween20 0.05%), permeabilized with 0.2% Triton X-100 in PBS for 10 min, washed again three times with PBST.5, followed by a 30 min incubation in 70% EtOH at room temperature. Coverslips were then dehydrated with subsequent washes with 80%, 90% and 100% EtOH for 5 min each and briefly dried.

Xist probes were generated from an Xist cDNA plasmid (∼18 kb) using a nick translation kit (Abbott) and Cy5-dUTP (Enzo) following the manufacturers protocol with incubation for 16 h at 15°C. Cy5 labelled Xist probes, 3µl per hybridization, were ethanol precipitated with 1/3 volume of 10 mg/ml salmon sperm DNA, 1/10 volume of 3 M NaOAc and 3 volumes of 100% EtOH. After spinning and washing with 70% EtOH, the pellet was air dried and resuspended in deionized formamide (Sigma). Probes resuspended in formamide were added to the hybridisation mixture to a final concentration of 2xSSC buffer, 50% Formamide (containing probes), 5% Dextran sulphate, 1 mg/ml BSA, 2 mM Vanadyl Ribonucleoside Complex.

Coverslips were hybridized with 30µl probe/ hybridization buffer mix that was denatured at 75°C for 5 min before briefly being chilled on ice. Hybridization was carried out in a humid chamber overnight at 37°C. The next day, coverslips were washed three times with prewarmed 50% formamide/2x SSC at 42°C for 5 min each, and subsequently three times with 2x SSC at 42°C for 5 min each. Coverslips were rinsed with water to remove excess salts and mounted with Vectashield antifade mounting medium containing DAPI (Vector Labs) on Superfrost Plus microscopy slides (VWR). Slides were dried, sealed using clear nail polish, and cleaned for imaging.

For immunofluorescence, cells were cultured on gelatin-coated No. 1.5H precision coverslips (170 ± 5 µm; Marienfeld Superior) and treated with dox for 24h. Coverslips were washed once with PBS prior to fixation in 3.7% formaldehyde for 10 mins. Coverslips were washed 3x with PBST.5, permeabilized with 0.2% Triton X-100 in PBS for 10 min and washed again 3x with PBST.5. Coverslips were blocked in 4% BSA diluted in PBST.5 for 30 mins. A primary antibody against H2AK119Ub (CST - #8240) was used at a 1:1000 dilution in 4% BSA and incubated overnight at 4°C. Coverslips were washed 3x in PBST.5 and incubated with an AlexaFluor647 conjugated goat anti-mouse IgG secondary antibody (Invitrogen - 1:2000) for 1h in 4% BSA. Coverslips were washed 3x in PBST.5 followed by washes in PBS and ddH20 prior to mounting in Vectashield with DAPI (Vector labs).

### ChrRNA-seq

Chromatin-associated RNA (ChrRNA) samples were prepared as previously described^16^. Briefly, 5 x106 cells per condition were collected washed once in PBS, pelleted and snap frozen on dry ice. Cell pellets were resuspended in hypotonic buffer + NP-40 to isolate nuclei with cytoplasmic debris separated using an underlayer of buffer containing a high concentration of sucrose. Nuclei were pelleted and resuspended in a buffer containing high concentrations of NP-40 and Urea to lyse nuclei. Chromatin was pelleted and resuspended in a high salt buffer and digested with Turbo DNaseI for 30 mins (Invitrogen) followed by proteinase K digestion (Roche - 1mg/ml). After digestion, RNA was isolated using Trizol reagent (Thermo fisher) and precipitated using isopropanol and subject to a second round of DNaseI treatment. ChrRNA purification was then carried out using an RNA clean-up kit (Zymo). The integrity of ChrRNA was analysed using a Bioanalyser and RNApico kits (agilent) to confirm the presence of nascent transcripts including pre-rRNA products. ChrRNA-Seq libraries were then prepared using the Illumina TruSeq RNA kit using rRNA depletion beads (Vazyme) and a total of 500 ng of ChrRNA and following the manufacturers protocols. Library concentrations were quantified using Qubit dsDNA high sensitivity reagents and library sizes were quantified using a Bioanalyser and high sensitivity DNA kits. Libraries were then pooled and sequenced either using NextSeq500 or NovaSeq platforms. Allelic analysis of Xi-specific gene expression and Xist expression was conducted as previously described^24^.

### Western blot

For whole-cell extracts mESCs were lysed directly in 96-well plates using 150 µl of 2X SMASH buffer (33 mM Tris-HCl (pH 6.8), 11% Glycerol, 40 mg/ml SDS, 200 mg/ml bromophenol blue and 10% β-mercaptoethanol in nuclease-free water). Samples were incubated for 1h at room temp prior to 3 freeze thaw cycles on dry ice to reduce the viscosity and DNA content of the samples. Samples were then run on 8% SDS polyacrylamide gels for 1h at 200V, transferred onto nitrocellulose membranes using a Biorad TurboBlot using the mixed MW settings. Blots were blocked in 5% milk in TBST prior to incubation with primary antibodies. Western blots for dT-HNRNPK used antibodies against HNRNPK (CST #4675 - 1:1000) and α-Tubulin (CST - #2144 - 1:1000) as a loading control. Blots for HNRNPU-dT used antibodies against HNRNPU (made in house – 1:6000) and METTL3 (abcam - ab195352 - 1:6000) as a loading control. Primary antibodies were incubated at 4°C overnight and washed thoroughly with TBST prior to secondary antibody incubation with anti-Rabbit-HRP (Amersham - NA934 - 1:2000). Blots were then washed 3x with TBST, once with PBS and incubated with Clarity ECL reagent (Biorad) prior to developing on radiographic film (Amersham).

### Live-cell imaging

Cells were seeded in gelatin-coated Ibidi 8-well 1.5H polymer imaging slides. For short-term imaging experiments cells were treated with dox and dTAG-13 as required, washed once with PBS before being imaged in live-cell imaging media (see above). If Halo imaging was required, cells were labelled with Halo-JFX650 (Promega) prior to imaging following the manufacturer’s instructions. For long-term imaging experiments, cells were treated with 1µM Trolox (Sigma) 24h before imaging to reduce the effects of phototoxicity. Cells were treated with dTAG-13, if required, 2h prior to imaging and dox induction at was carried out in 8-well imaging slides whilst on the microscope and imaged directly after induction.

### Fixed-cell imaging

To accurately determine Xist spreading changes during the cell-cycle, cells were cultured in 8-well imaging dishes (ibidi), labelled with Halo-JFX650 to image Halo-PCNA and fixed in 4% paraformaldehyde for 10 mins at room temp. Cells were washed 3x with PBST.5, ddH2O and covered with Vectashield containing DAPI directly in the imaging slide. Slides were incubated overnight at 4°C prior to imaging.

### Microscopy

Ibidi imaging slides for both fixed and live-cell Nuc-BRITE samples were imaged using a Visitech iSIM super resolution unit built around an Olympus IX83 microscope. The iSIM has 4 diode lasers 405, 488, 561 and 640 nm (10 mW each) that run through the iSIM super-res confocal array scanner and images collected on a dual camera set-up housing two Hamamatsu Orca Quest cameras. Fluorescence was filtered using specific pairs of filters for each pair of channels: 488/561, 488/640, 405/640. Fast super-resolution dual-colour imaging used a 1.5 N.A. 100X PlanApo objective, 63 ms exposure and 80% laser power for 200 frames. Long-term imaging used a 1.4 N.A. 60X PlanApoTIRF objective (Olympus), 63ms exposure and 12% laser power (488/561) over multiple Z-sections (0.3 µm) every 2 mins for 2h. The dual-camera set-up was aligned across all 4 channels using X, Y and Z tilt adjustments and a TetraSpek bead slide (Thermo Fisher) to compensate for any chromatic aberrations. Live-cell LacI domains and fixed IF samples were imaged using an Olympus IX83 widefield microscope using CooLED illumination at 365, 490, 580 and 635 nm through a 1.35 N.A. 60X PlanApo objective, filtered using fixed filter sets (DAPI, GFP, TRITC and Cy5) and images were collected on a Prime 95B sCMOS camera (Photometrics). Images were acquired as Z-stacks using 10ms (DAPI) or 200ms (GFP/TRITC/Cy5) exposure time and 5% (DAPI) and 30% (GFP/TRITC/Cy5) illumination.

### Image analysis

Analysis of Xist RNA intensities, dynamics and spreading was undertaken using ImageJ, the Trackmate plugin^31^ and custom MATLAB scripts. In short, Xist clouds were cropped, batch processed using the Trackmate batcher plugin which exported trajectory and foci information which was subsequently imported in MATLAB for quantification of intensities, diffusive behaviours and spreading on a cell-by-cell basis. More specifically, to quantify intensities, cropped images were batch processed with the following Trackmate settings to detect single particles and reduce false positives from neighbouring or out-of-focus foci: Laplacian of Gaussian detector, Foci diameter – 200 nm, Quality – 15, Contrast – 0.03, LAP tracker – Frame linking 200 nm, Gap closing 200 nm, Number of spots in track – 5. Only spots that were allocated to trajectories were analysed and .csv files were imported into MATLAB for quantification and data plotted as histograms in Igor Pro 9 (Wavemetrics). Foci intensities were normalised to foci observed in mESCs transfected with a plasmid expressing a MS2v7 x24 array alone. These data were acquired on the same day with identical settings to robustly normalise the Xist data by simply subtracting the background intensity outside of cells and dividing by the average MS2v7 x24 intensity. Signal-to-noise ratios were quantified by dividing the mean foci intensity by the mean background intensity adjacent to the foci, for example, nuclear background was used for Xist and cytoplasmic background for MS2v7 x24.

For quantification of diffusive behaviour and trajectory information, images were batched processed using Trackmate and the following settings to build trajectories in a robust manner: Laplacian of Gaussian detector, Foci diameter – 200 nm, Quality – 20, Contrast – 0.05, LAP tracker – Frame linking 200 nm, Gap closing 200 nm, Number of spots in track – 5. Slight alterations to intensity and gap closing distances had to be made to accurately quantify Mitotic, SPEN KO, SPEN ΔRRM and hnU-dT + dTAG-13 datasets due to reduced foci intensities, increased distances between foci and increases in mobilities: Laplacian of Gaussian detector, Foci diameter – 200 nm, Quality – 15, Contrast – 0.05, LAP tracker – Frame linking 200 nm, Gap closing 600 nm, Number of spots in track – 5. These global settings to analyse imaging datasets were validated by processing random samples of images by eye to ensure particles were tracked accurately with minimal errors. Information on trajectories such as track length and trajectories per cell were quantified from the tracks .csv output of Trackmate. Diffusion coefficients were calculated using the msdanalyser^32^ suite of processes in MATLAB. Briefly, cropped images were batch processed in Trackmate batcher and simple .xml files were exported and subsequently imported as concatenated cell arrays into MATLAB. Cell arrays were then processed individually in msdanalyser to correct for drift on a cell-by-cell basis, the first 25% of MSD curves were fitted to calculate a diffusion coefficient for each trajectory. Fits which had an r^2^ value less than 0.6 were excluded and all diffusion coefficients (D) were compiled, corrected for 75 ms per frame and plotted as log_10_D values in Igor Pro 9.

Analysis of Xist RNA spread in fixed samples was quantified in a similar way as with the live-cell imaging described previously but analysing foci in 3D rather than 2D over time. Cropped images were batch processed in Trackmate using the following settings: Differences of Gaussian detector (DoG), Foci diameter – 200 nm, Quality – 15, Contrast – 0.05 and exported as ‘all-spots.csv’ files. Foci positions as X, Y and Z co-ordinates were used to calculate numbers of foci per cell and Xist cloud volumes by calculating a convex hull volume in MATLAB using foci co-ordinates.

Long-term Xist spreading was quantified by tracking particles from 2D maximum projections of 3D acquired images on a cell-by-cell basis over time using the following Trackmate settings: Laplacian of Gaussian detector, Foci diameter – 200 nm, Quality – 20, NND tracker – Frame linking 2 µm and Number of spots in track – 3. MATLAB was used to quantify the intensity of each focus over time and calculate the area of the Xist cloud using a convex hull from the X, Y co-ordinates. These data were split by frame into groups of foci intensities over time which allowed for the quantification of the maximum and minimum intensities which represented the putative nascent transcription site and spread Xist foci. Plotting histograms of foci intensities allowed for the normalisation of number of molecules observed per nascent RNA in both wild-type and mutant contexts.

In MATLAB, interactions between Xist foci were analysed from the fast acquisition data to calculate the mean X, Y and T positions of all tracks, a nearest neighbour search of these mean trajectory positions was obtained and neighbouring trajectories were paired ensuring each pair was unique and paired with one another. Paired trajectories with an average NND greater than 400 nm apart were excluded from further analysis and euclidean distances between the remaining pairs were calculate over time for each nearest neighbour pair independently. These NND vs time plots were fitted with a maximum likelihood estimation model previously used for detecting changes in fluorescent intensity trajectories and from these fits, significant changes in distance between pairs (ΔNND) and fractional durations of nearest neighbors less than 400 nm were quantified. All MATLAB scripts can be found on our github page: https://github.com/adamcawte/Xist_MATLAB_Code

## Acknowledgments

We would like to thank members of the Brockdorff lab, Alexander Borodavka and Stephan Uphoff for critical reading of this manuscript. We thank Luca Georgetti (FMI, Switzerland) for plasmids for the LacO array and LacI** protein.

## Funding

This work was funded by grants to NB from the Wellcome Trust (215513/Z/19/Z) and UKRI (EP/Y029062/1). Imaging was performed at the Micron Oxford Advanced Bioimaging Unit funded by a Wellcome Trust Strategic Award (091911 and 107457/Z/15/Z).

## Author contributions

N.B. and A.D.C conceived experiments. N.B. and A.D.C wrote the manuscript. A.D.C generated and validated cell lines, conducted imaging experiments and image analysis. G.W. ran computational analysis of ChrRNA-seq data. F.C, M.A and H.C generated plasmids and helped facilitate the production of mutant cell lines.

## Competing interests

The authors declare no competing interests.

## Data, code, and materials availability

All data, code, and materials used in this study are available upon request. ChrRNA-seq data is available through the Gene Expression Omnibus under the accession code - GSE331175. MATLAB data analysis pipelines for Xist SPT data are available on github: https://github.com/adamcawte/Xist_MATLAB_Code.git

## Supplementary Materials

Supplementary Notes 1 - 2

Supplementary Figures 1 - 5

Supplementary Table 1

Supplementary Video legends 1 - 7

## Notes

### Competing Interest Statement

The authors have declared no competing interest.

