## Supplementary Information for "A contact-mediated transfer mechanism directs *in cis* localization and spread of Xist RNA"

**This file includes:**

Supplementary Notes 1 - 2
Supplementary Figures 1 - 5
Supplementary Table 1
Supplementary Video Legends 1 – 7
Supplementary References

### Supplementary Notes

Supplementary Note 1: Quantification of Xist foci interactions

To quantify the interactions made by neighbouring Xist molecules we focused our analysis on the distances neighbouring molecules either converge or move away from one another, as well as the duration of the trajectory that they remain close together. For this we calculated the mean position in X, Y and T for each Xist trajectory in a cloud and paired nearest neighbour trajectories. If these neighbouring trajectories were correctly paired with each other, unique, and less than 400 nm apart on average, they were kept for further analysis, with other trajectories being discarded. Nearest neighbour distance (NND) trajectories were calculated between pairs and the profiles were fitted with a maximum likelihood estimation[1,2] to quantify significant changes in NND (ΔNND) and the time molecules spent less than 400 nm apart as a fraction of trajectory length (Fig. 3c). We observed a median ΔNND value of ~ 150 nm for wild-type cells (Fig. 3d), suggesting small, confined movements consistent with chromatin associated dynamics. The duration of foci interactions in wild-type cells, shows a bi-modal distribution with most pairs spending either 100% of the trajectory less than 400 nm and the other main population at ~10% with a median value of ~ 45% (Fig. 3e). More importantly, these data provide a way to assess the variability in Xist-Xist interactions over time.

Supplementary Note 2: Investigation of HNRNPK involvement in Xist RNA spreading

Imaging of Xist and PCNA during different stages of S-phase in Xist-ΔPID or dT-hnK + dTAG-13 lines indicated that Xi replication occurs in early S-phase (Fig 2f-h and Supplementary Fig 5a), contrasting with our analysis of wild-type cells and prior work[3] demonstrating Xi replication is in mid S-phase. To quantify this observation, we took the ratio of Halo-PCNA over the Xi domain (as marked by the dense region of Xist signal) relative to the Halo-PCNA signal in the nucleoplasm in early S-phase cells. We observed that Xist-ΔPID or dT-hnK + dTAG-13 lines cell lines show an increased PCNA Xi:Nuc ratio closer to 1 showing that Xi replication indeed starts earlier in S-phase (Supplementary Fig 5b). Absence of PRC1-mediated H2AK119ub, irrespective of Xist cloud size, was also confirmed quantitatively in both Xist-ΔPID and dT-hnK +dTAG-13 cell lines (Supplementary Fig 5c and d).

A prior study reported that compaction of Xi territories is markedly reduced as a consequence of Xist B-repeat deletion (approximately equivalent to Xist-ΔPID used herein)[4]. We observed low intensity accumulation of LacI-tdTomato signal in wild-type Nuc-BRITE mESCs surrounding the LacOx100 foci specifically upon dox induction for 24h (Fig 1b, Supplementary Fig 5e and Supplementary Video 1). LacI is a constitutive DNA binding protein which slides to find its target LacO sequence where residency is sufficiently high[5], and accordingly we interpret the low intensity LacI-tdTomato signal as non-specific association across the compacted Xi territory. Interestingly, this low intensity signal was not observed upon Xist induction in the Xist-ΔPID and or dT-hnK + dTAG-13 lines (Supplementary Fig 5e and f). We interpret this observation as indicating reduced compaction of Xi, likely attributable to the absence of Polycomb-mediated histone modifications.

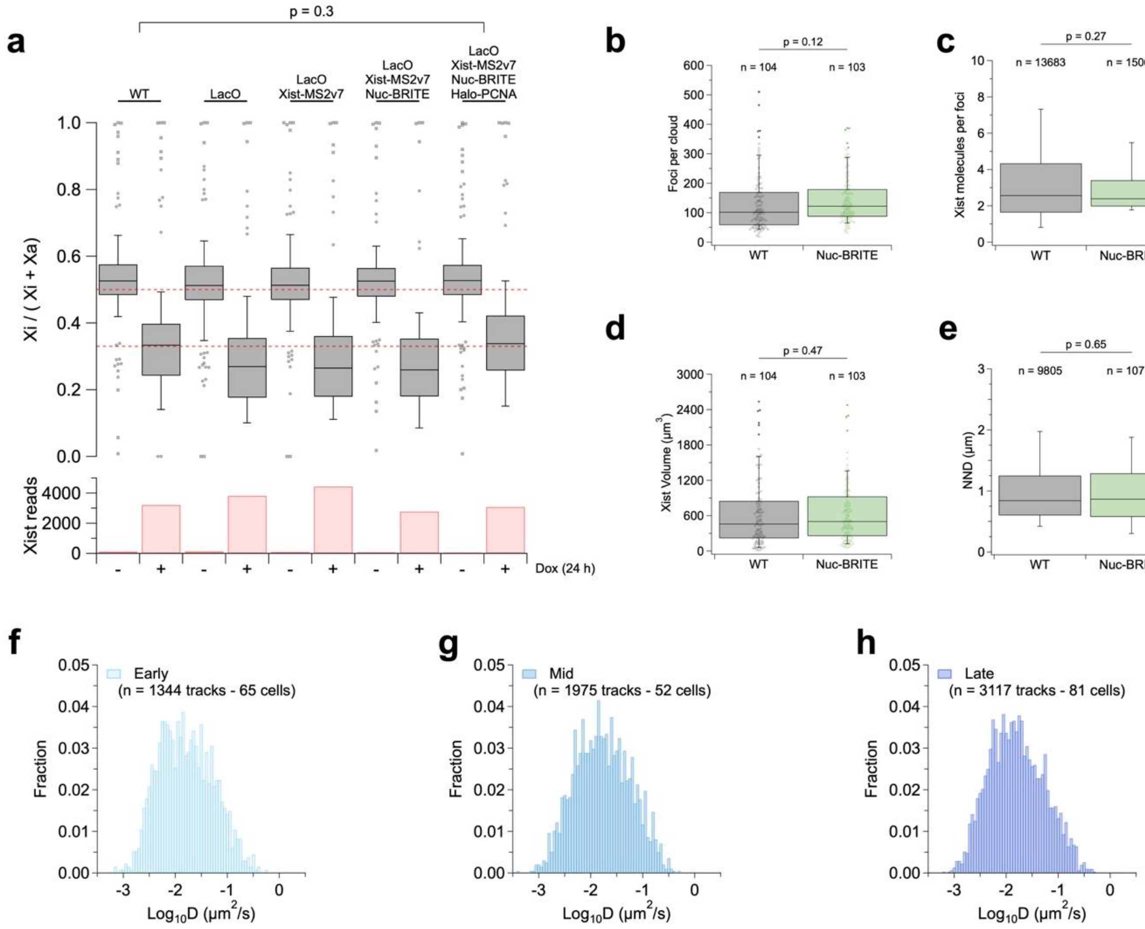


**Supplementary Fig. 1**

(a) X chromosome silencing plot showing the Xi gene expression ratio as Xi/(Xi+Xa) for each modification to generate Xist-Nuc-BRITE cells with or without dox induction for 24h. Red dashed lines depict silencing ratio in the absence of dox (0.5) and following dox induction (~0.33) for wild-type cells (WT). Below, shows bar graph of Xist read count as RPKM in no dox and +dox conditions for each cell line. The number of X-linked genes analysed for each sample were 257, 256, 256, 243 and 279 for WT, LacO, LacO/MS2v7, LacO/MS2v7/Nuc-BRITE and LacO/MS2v7/Nuc-BRITE/Halo-PCNA respectively. Quantification of RNA-FISH foci in wild-type (WT) and Xist-Nuc-BRITE

(Nuc-BRITE) cells for foci number (b), number of Xist molecules per focus (c), Xist cloud volume (d) and Xist nearest-neighbour distance (NND) (e). N values represent number of cells (b and d), number of foci (c) and number of unique NNDs (e). (f - h) $Log_{10}$ diffusion coefficient histograms for early, mid and late S-phase stages. N displayed as number of tracks and number of cells analysed. P-values shown calculated from unpaired single tailed t-tests.

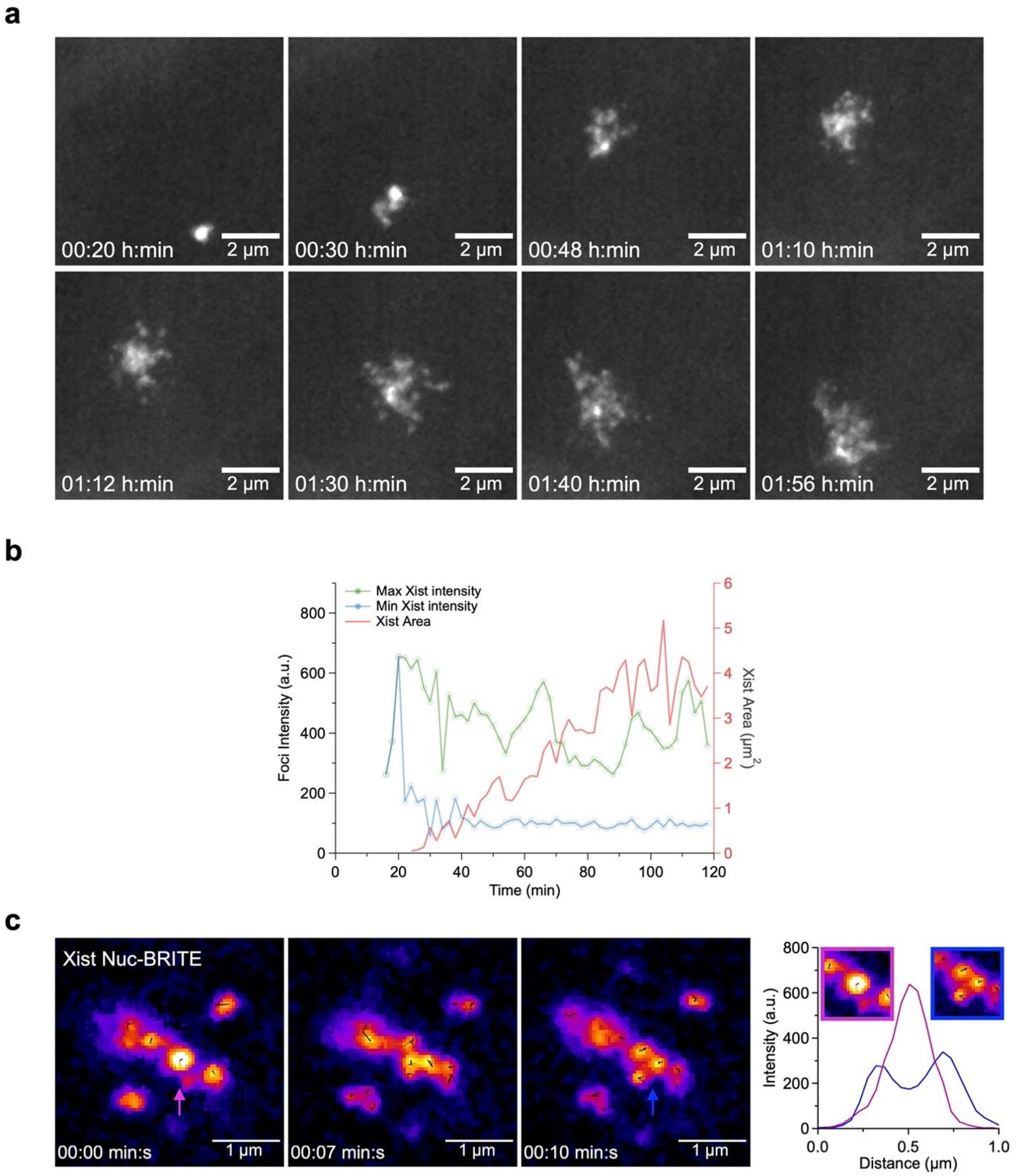


**Supplementary Fig. 2**

(a) Individual frames from 2h time course of Xist expression and initial spreading events (Supplementary Video 4) showing bright nascent transcription foci and dimmer spreading Xist foci. (b) Foci intensity plots for the minimum (blue) and maximum (green) Xist intensities over time as well as the calculated Xist area (red) for the Xist spreading depicted in (a) and Supplementary Video 4. (c) Individual frames of Xist RNA-RNA interactions following 2h of dox induction (Supplementary Video 5) showing merged Xist foci (magenta arrow and intensity profile) and split Xist foci (blue arrow and intensity profile) dynamically interact in the early stages of Xist spreading.

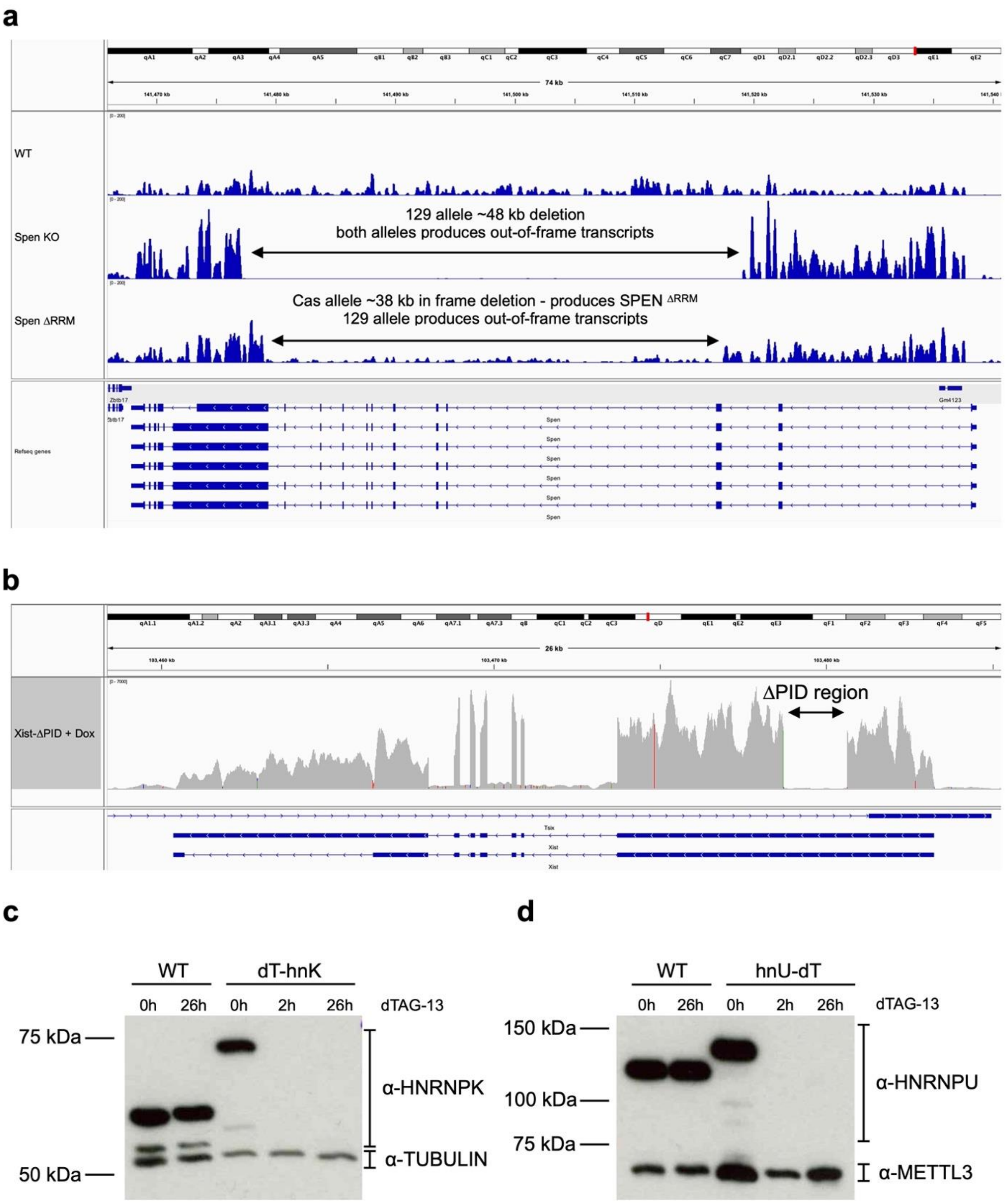


**Supplementary Fig. 3**

(a) ChrRNA-Seq profile for the *Spen* locus in wild-type Nuc-BRITE (WT), SPEN $^{KO}$ and SPEN $^{\Delta RRM}$ cells showing the 129 allele ~48kb out-of-frame deletion in the SPEN $^{KO}$ line and the Cast allele ~38kb in frame deletion to produce mono-allelic expression of a SPEN $^{\Delta RRM}$ product. (b) ChrRNA-Seq profile of the Xist locus following Xist induction in the Xist-∆PID cells showing the deletion of the ∆PID region on the 129 Xi allele. (c) and (d) show western blots of wild-type Nuc-BRITE cells compared to dT-hnK and hnU-dT cells treated either with 2h or 26h of dTAG-13 as annotated. Loading controls either used anti-α-TUBULIN or anti-METTL3 antibodies.

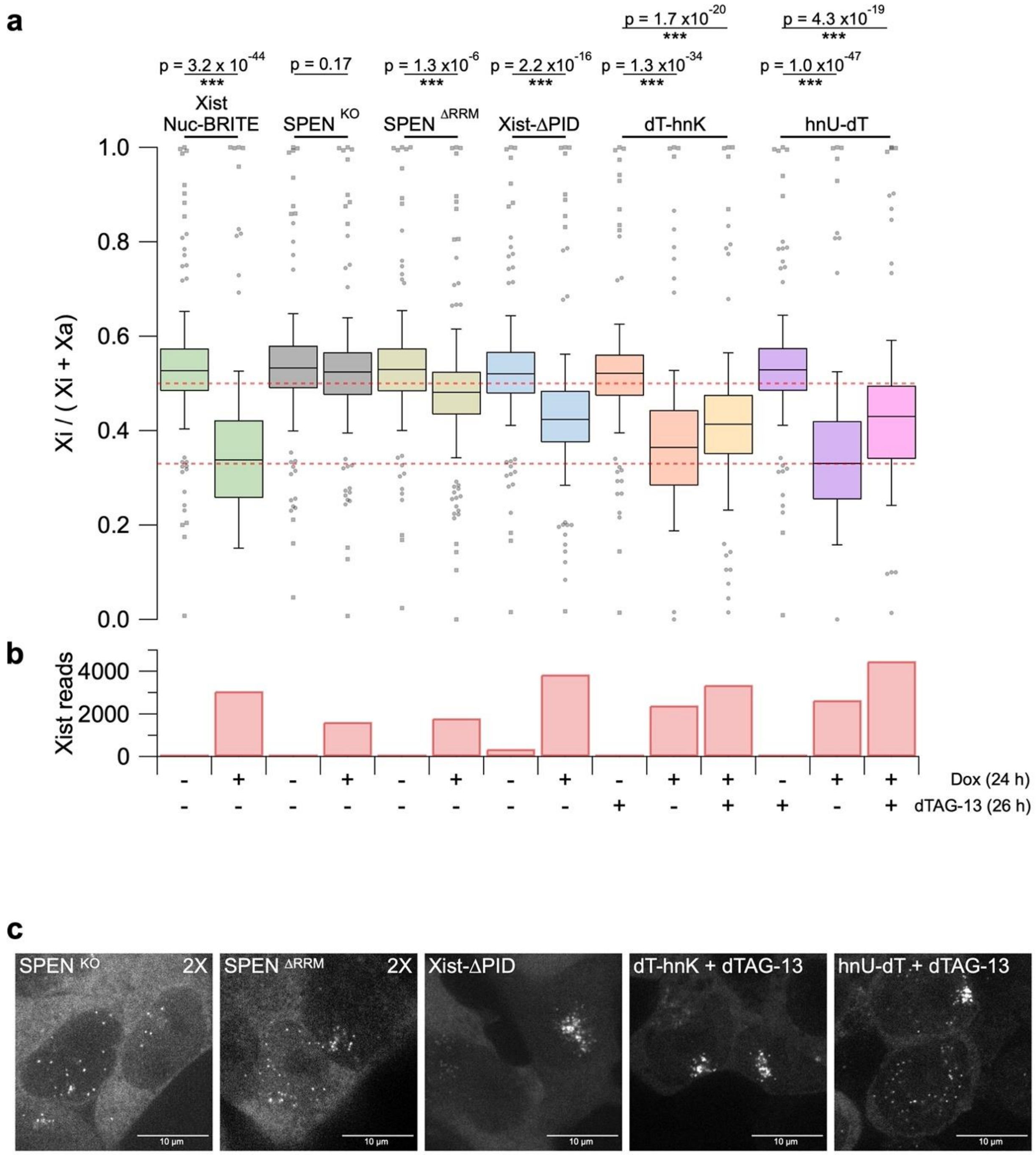


**Supplementary Fig. 4**

(a) X chromosome silencing plot showing the Xi gene expression ratio as Xi/(Xi+Xa) for wild-type Nuc-BRITE cells and each mutant line with and without dox or dTAG-13 where appropriate. Red dashed lines depict expected silencing ratio in the absence of dox

(0.5) and following dox induction (~0.33). (b) Shows bar graph of Xist read count as RPKM in no dox, +dox and +dox/+dTAG-13 conditions for each cell line where appropriate. Statistical analysis used a single-tailed t-test to compare the differences between conditions. All samples compared the same 279 X-linked genes which met the threshold for significant expression levels. (c) Representative images of Xist RNA spread in different mutant cell line conditions. SPEN $^{KO}$ and SPEN $^{\Delta RRM}$ are shown at 2X normal contrast to better visualize low intensity foci. Live-cell imaging data for these cells can be found in Supplementary Video 6.

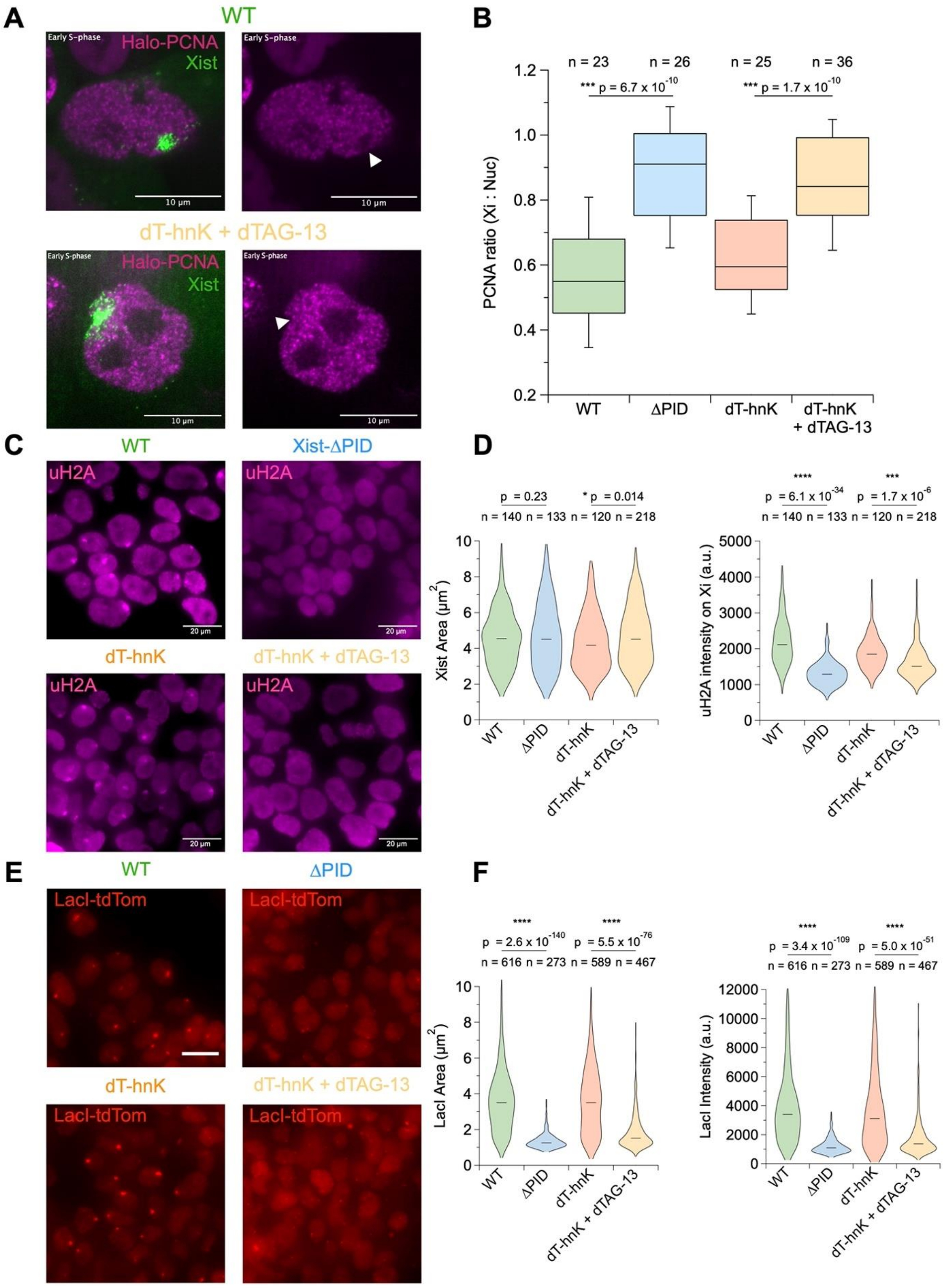

A
WT
Early S-phase
Halo-PCNA
Xist
10 µm
dT-hnK + dTAG-13
B
n = 23
n = 26
n = 25
n = 36
*** p = 6.7 x 10^-10
*** p = 1.7 x 10^-10
PCNA ratio (Xi : Nuc)
WT
ΔPID
dT-hnK
dT-hnK + dTAG-13
C
Xist-ΔPID
uH2A
20 µm
D
p = 0.23
* p = 0.014
n = 140 n = 133 n = 120 n = 218
Xist Area (µm²)
**** p = 6.1 x 10^-34
*** p = 1.7 x 10^-6
uH2A intensity on Xi (a.u.)
E
LacI-tdTom
F
**** p = 2.6 x 10^-140
**** p = 5.5 x 10^-76
n = 616 n = 273 n = 589 n = 467
LacI Area (µm²)
**** p = 3.4 x 10^-109
**** p = 5.0 x 10^-51
LacI Intensity (a.u.)


114

**Supplementary Fig. 5**

(a) Representative images for Halo-PCNA signal in early S-phase cells in WT and dT-hnK + dTAG-13 conditions. Arrows show Xi domain with low or high PCNA signal in WT vs dT-hnK + dTAG-13 respectively. (b) Quantification of PCNA Xi : Nucleoplasm ratio in wild-type Nuc-BRITE (WT), Xist-ΔPID (ΔPID), dT-hnK and dT-hnK + dTAG-13 cells. N represents the number of cells analyzed and statistical analysis using a single-tailed t-test. (c) Representative images of H2AK119ub (uH2A) antibody staining in WT, ΔPID, dT-hnK and dT-hnK + dTAG-13 cells. (d) Quantification of Xi domain area (using Xist-Nuc-BRITE signal) and the mean uH2A intensity within the detected Xi domain. N represents the number of cells analyzed and statistical analysis using a single-tailed t-test. (e) Representative images of LacI-tdTom signal in WT, ΔPID, dT-hnK and dT-hnK + dTAG-13 cells. (f) Quantification of LacI area and the mean LacI intensity for each detected domain. N represents the number of cells analyzed and statistical analysis using a single-tailed t-test.

**Supplementary Table 1**

**Table of oligos used for CRISPR sgRNAs and PCR screening.**

Lower case **g** in sgRNA denotes additional base required for PolIII transcription. SPEN screening primer (red) used all three in combination to detect partial loss of exon 3.

| Target | sgRNA seq | Screening primer F | Screening primer R |
|---|---|---|---|
| **Rosa26 locus** | **g**CGCCCATCTTCTAGAAAGAC | CCGCCTCCTCCGAGGACAACAACA | ACAAGCTCACAAGACCTTAGG |
| **Tsix – LacO insert** | **g**CTGGGAACACGAGGAGCCCA | ATGCTCTCATAAACAACGTGGAAGG | TGTTATTACACGTTGCTGTCTCTTCTATC |
| **Xist – MS2v7 insert** | **g**CATACGTAGTTCCCCGCTCT | TCTGGTTGCTATTTCTCTTAGCTTTTC | GGAATATGCAACTAAGACCATGAACCC |
| **SPEN 5'** | **g**CGGACAAGACATTACGATC | CAGAAAGAGGCGAGGCGTAAAG | ATAGGCACTATGTTCATAAGCACGC |
| **SPEN 3'** | GGGGTGTCTCCTGCGCATT | N/A | GTCGTCCCTGTAGATATCCCTGTG |
| **∆PID** | **g**TTGAGGTTCTATACCAGTTC | GTTTCGCTTGGTACTTCACG | GCAAGGGTAGTATTAGGACC |
| **HNRNPK – Nterm** | **g**CTTCTCACCAAATTCACCAT | N/A | N/A |
| **HNRNPU - Cterm** | **g**AAACAGTCGACTTCTTGTGA | ggcagaagtagaggggaagg | gccttcccgttttgtcagag |

**Supplementary Video 1**

Fast frame rate movies (75ms) acquired on the VisiTech iSIM of Xist-Nuc-BRITE and LacO-tdTomato signals with 24h of dox induction. Scale bars are 10 µm.

**Supplementary Video 2**

Fast frame rate movie (75ms) acquired on the VisiTech iSIM of Xist-Nuc-BRITE cells transfected with pCAG-MS2v7x24-pA plasmid for a visual comparison of monomeric mRNA molecules in the cytosol and Xist multimeric accumulation in the nucleus. Cells expressing MS2v7x24 annotated with blue arrows and cells not expressing MS2v7x24 annotated with green arrows. Quantification of number of RNA molecules per focus shown as a histogram normalized to the mean of MS2v7x24 intensities. Data taken from fast framerate imaging where the number of foci analysed is represented as n taken from 10 and 50 cells for MS2v7x24 and Xist-Nuc-BRITE samples respectively.

**Supplementary Video 3**

Fast frame rate movie (75ms) of Xist-Nuc-BRITE cells with 24h dox induction acquired on the VisiTech iSIM showing interactions of Xist foci. Foci were tracked using trackmate and trajectories are shown in a uniform colour (black) overlaid on a fire LUT set in FIJI. Scale bar is 1 µm.

**Supplementary Video 4**

Long-term imaging of Xist RNA spreading over 2h at a 2 min frame rate. Nascent Xist production can be observed as a bright focus followed by initial Xist spreading events creating dimmer foci adjacent to the nascent Xist foci. Plotting the minimum and maximum Xist foci intensities allows for the detection of nascent Xist accumulation and individual Xist foci spreading to new sites on the Xi. This data is plotted along with the Xist area for the adjacent movie to show how we compute nascent Xist RNA intensities.

**Supplementary Video 5**

Fast frame rate movie (75ms) of Xist-Nuc-BRITE cells with 2h dox induction acquired on the VisiTech iSIM showing Xist foci interacting. Foci were tracked using trackmate and trajectories are shown in a uniform colour (black) overlaid on a fire LUT set in FIJI. Scale bar is 1 µm.

**Supplementary Video 6**

Fast frame rate movies (75ms) of Xist spreading phenotypes in mutant cell lines for SPEN $^{KO}$, SPEN $^{\Delta RRM}$, Xist-ΔPID, dT-hnK + dTAG-13 and hnU-dT +dTAG-13 with 24h dox induction acquired on the VisiTech iSIM. SPEN $^{KO}$ and SPEN $^{\Delta RRM}$ are set to 2x the contrast to better visualize the relatively low intensity foci that were observed. All other cells are set to the same contrast settings. Scale bar is 10 µm.

**Supplementary Video 7**

Fast frame rate movies (75ms) for Xist-Nuc-BRITE, SPEN $^{KO}$, dT-hnK + dTAG-13 and hnU-dT + dTAG-13 cells with 24h dox induction acquired on the VisiTech iSIM showing the behaviour of interacting Xist foci. Foci were tracked using trackmate and trajectories are shown in a uniform colour (black) overlaid on a fire LUT set in FIJI.
